# Migration through a major Andean ecogeographic disruption as a driver of genotypic and phenotypic diversity in a wild tomato species

**DOI:** 10.1101/2020.09.09.289744

**Authors:** Jacob B. Landis, Christopher M. Miller, Amanda K. Broz, Alexandra A. Bennett, Noelia Carrasquilla-Garcia, Douglas R. Cook, Robert L. Last, Patricia A. Bedinger, Gaurav D. Moghe

## Abstract

The large number of species on our planet arises from the phenotypic variation and reproductive isolation occurring at the population level. In this study, we sought to understand the origins of such population-level variation in defensive acylsugar chemistry and mating systems in *Solanum habrochaites* – a wild tomato species found in diverse Andean habitats in Ecuador and Peru. Using Restriction-Associated-Digestion Sequencing (RAD-seq) of 50 *S. habrochaites* accessions, we identified eight population clusters generated via isolation and hybridization dynamics of 4-6 ancestral populations. Estimation of heterozygosity, fixation index, isolation by distance, and migration probabilities, allowed identification of multiple barriers to gene flow leading to the establishment of extant populations. One major barrier is the Amotape-Huancabamba Zone (AHZ) – a geographical feature in the Andes with high endemism, where the mountainous range breaks up into isolated microhabitats. The AHZ was associated with emergence of alleles for novel reproductive and acylsugar phenotypes. These alleles led to the evolution of self-compatibility in the northern populations, where alleles for novel defense-related enzyme variants were also found to be fixed. We identified geographical distance as a major force causing population differentiation in the central/southern part of the range, where *S. habrochaites* was also inferred to have originated. Findings presented here highlight the role of the diverse ecogeography of Peru and Ecuador in generating new, reproductively isolated populations, and enhance our understanding of the microevolutionary processes that lay a path to speciation.

## Introduction

Heritable phenotypic variation, adaptation and reproductive isolation between populations are recognized as the primary drivers of speciation in evolutionary theory (Darwin, 1859; Reznick & Ricklefs, 2009; Harvey *et al*., 2019). Thus, studying how new traits arise in populations is crucial to our understanding of the emergence of biological diversity. The advent of next-generation sequencing technologies allows integration of phylogenetic analysis and mechanistic studies of trait variation across populations, helping improve our understanding of microevolution (Harvey *et al*., 2019). In this study, we utilized Restriction Associated Digestion Sequencing (RAD-seq) (Miller *et al*., 2007; Baird *et al*., 2008) in the wild tomato *Solanum habrochaites* – a species well-studied for its population-level diversity – to assess demography, defense metabolites and reproductive traits in an integrative manner.

*Solanum habrochaites* (Knapp & Spooner, 1999) is a phenotypically diverse species with a range from the upper reaches of the Atacama desert in southern Peru to the tropical forests of central Ecuador. Growing along the western Andes, this species is generally found 1000-3000 meters above sea level (masl) but extends down to sea level in central Ecuador. This diversity in distribution and habitat may be at least partially responsible for the observed phenotypic diversity in this species, which is described below.

Previous studies (Gonzales-Vigil *et al*., 2012; Kim *et al*., 2012; Schilmiller *et al*., 2015; Fan *et al*., 2017) have demonstrated substantial population variation in two trichome-localized compound classes – acylsugars and terpenes – that are important for defense against herbivores (Weinhold & Baldwin, 2011; Leckie *et al*., 2016). For example, *S. habrochaites* accessions were grouped into two chemotypic superclusters based on their acylsugar profiles – a “northern” supercluster that failed to add an acetyl (C2) group to the sucrose R2 position in acylsugars, and a “southern” supercluster that retained this activity (Kim *et al*., 2012). This loss of C2 addition was a result of inactivation of acylsugar acyltransferase 4 (ASAT4), the final enzyme in the *Solanum* acylsugar biosynthetic pathway. This inactivation occurred via three different mechanisms – loss of gene expression, frameshift mutation and likely gene loss in different accessions. Using the same individuals sampled in this project, another study demonstrated differential acylation between northern and southern accessions on the furanose ring of the acylsugar, which could be traced back to gene duplication, divergence and loss in ASAT3, an upstream enzyme in the pathway (Schilmiller *et al*., 2015). However, demographic processes that influenced this evolution of acylsugar profiles are not known.

*S. habrochaites* is also an attractive system for the study of reproductive trait evolution, with extensive diversity both in mating system and in reproductive barriers that affect gene flow between populations and between *S. habrochaites* and other tomato clade species. *S. habrochaites* is predominantly an obligate outcrossing species due to gametophytic S-RNase-based self-incompatibility (SI) (Mutschler & Liedl, 1994; Peralta *et al*., 2008; Bedinger *et al*., 2011). In this type of SI, the *S*-locus encodes pistil-expressed S-RNases and pollen-expressed *S*-locus F-box proteins that determine the specificity of the SI interaction. In addition, other pistil-expressed (e.g., HT protein) and pollen-expressed (e.g. CUL1) factors that are not linked to the S-locus play a role in self pollen rejection [reviewed in (Bedinger *et al*., 2017)]. SI is widespread in flowering plants, and acts to preserve genetic diversity and diminish inbreeding depression (Stebbins, 1957; Lande & Schemske, 1985; Schemske & Lande, 1985; Takayama & Isogai, 2005; Igic *et al*., 2008). However, there may be a selective advantage for transitions to self-compatibility (SC) during the dispersal of species, since a single SC individual could conceivably colonize a novel environment in the absence of other individuals or pollinators (Baker, 1955, 1967; Stebbins, 1957; Pannell *et al*., 2015), which can set the stage for speciation (Allmon, 1992). SC *S. habrochaites* populations have arisen at the northern and southern species range margins (Martin, 1961; Rick *et al*., 1979). These marginal SC populations, located in Ecuador and southern Peru, represent independent SI-->SC transitions (Rick & Chetelat, 1991).

The populations at the northern species margin are of special interest, due to the diversity of reproductive barriers acting at the individual, population, and species levels (Martin, 1961; Broz *et al*., 2017b). In previous work, two distinct SC groups (SC-1 and SC-2) associated with different *S-RNase* alleles as well as differences in inter-population and interspecies crossing barriers were identified at the northern species range margin (Broz *et al*., 2017b). As *S. habrochaites* dispersed northward from its presumed site of origin in central Peru (Rick *et al*., 1979; Peralta *et al*., 2008; Pease *et al*., 2016), it traversed the Amotape-Huancabamba Zone (AHZ), a region of cordillera disruption near the Ecuador-Peru border that constitutes a barrier to species dispersal (Sillitoe, 1974; Weigend, 2004). The AHZ is bounded in the south by Río Chicama in Peru and in the north by Río Jubones in Ecuador (Weigend, 2002, 2004). A floristically diverse region called the Huancabamba Depression (HD) – coinciding with Río Huancabamba/Río Camaya/Río Marañon – is located in the central part of the AHZ in Peru (Weigend, 2002; Richter *et al*., 2009). With its highly variable microhabitats, the AHZ acts as a biodiversity hotspot for both plants and animals (Berry, 1982; Weigend, 2002), and may have influenced *S. habrochaites* evolution.

To determine how both SI-->SC transitions and acylsugar diversification occurred in the context of *S. habrochaites* range expansion, we first determined the species’ population structure using RAD-seq and studied patterns of and barriers to gene flow between different populations. We identified four independent SI-->SC transitions at the northern species margin associated with evolution of new acylsugar phenotypes. Our results revealed that alleles that eventually led to fixation of these novel phenotypes in Ecuador first emerged in the AHZ, during the northward migration of *S. habrochaites* from the Cajamarca region of Peru. We further observed the impact of geographical distance in central/southern Peru, producing locally isolated populations. This work underscores the critical role of ecogeography in shaping biological diversity.

## Materials and Methods

### Plant growth and sample collection for RAD-seq and biochemical analysis

At Michigan State University, 52 accessions of *S. habrochaites* and 4 accessions of *S. pennellii* (LA1941, LA1809, LA1674, LA0716) **(File S1**, plus LA2868, LA1978) were sterilized with 10% trisodium phosphate, germinated on moist filter paper and transferred to peat pots where they were grown for 2 weeks under 16:8 light:dark conditions at 25°C/16°C respectively. Up to four replicates of two-week old plants were then transferred to soil (2 Sure mix + 1/2 sand) where they were grown prior to their harvest for 2 more weeks under the same long day conditions with regular watering.

### Plant growth for reproductive phenotype analysis

At Colorado State University, seeds were sterilized according to recommendations of the TGRC (‘Tomato Genetics Resource Center’) and were planted into 4-inch pots containing ProMix-BX soil (Premier Tech Horticulture, Quakertown, PA, USA) with 16:8 light:dark conditions 26°C/18°C for 2 months. Plants were transplanted to outdoor agricultural fields at Colorado State University (May–September 2017) to obtain sufficient flowers for multiple crosses, and for collection of stylar tissue for immunoblotting analysis. For *S-RNase* allele analysis, plants were grown on a light shelf and a single young leaf was harvested from each plant for DNA preparation as previously described (Broz *et al*., 2017b).

### Library preparation and sequencing

Leaf tissue from one of the sampled individuals per accession was used for DNA extraction using the Qiagen DNeasy kit (Qiagen, Valencia, CA, USA). Integrity of DNA was verified as a single high molecular weight band on a 1% agarose gel. Biological replicates were obtained for two accessions (LA2098, LA2976 [2x]) and technical replicates for 17 accessions (LA1928, LA1731, LA1778, LA2976, LA1777 [2x], LA2975, LA1986, LA1352, LA2155, LA1737, LA2175, LA2098, LA1252, LA2105, LA2861, LA0407, LA1625 [2x]). Four accessions of *S. pennellii* were selected for outgroup analysis **(File S1)** – bringing the total number of RAD-seq samples to 78. One hundred ng of the extracted DNA was used for library preparation and sequencing in two Illumina HiSeq 2000 lanes, as described previously (von Wettberg *et al*., 2018).

Demultiplexed RAD-seq reads were deposited in NCBI Short Read Archive under the BioProject PRJNA623394.

### RAD-seq data processing

Overall, ∼198 million 100-bp single end reads were obtained after standard Illumina quality filtering. We first converted the FASTQ reads from Illumina 1.5 encoding to Sanger encoding using the seqret tool of the EMBOSS v6.5.0 package, trimmed the reads using FASTX toolkit v0.0.14 to a Phred score >20 and selected only 100b reads. Since the first base of all reads, which constituted part of the barcode, was ‘N’, it was trimmed away. The ∼187 million filtered reads were processed using the *process_radtags*.*pl* script in the Stacks software v2.3d (Rochette *et al*., 2019) with the following parameter settings *(-b barcodes_6b*.*tab -q -c -t 90 -E phred33 -D -w 0*.*20 -s 10 --inline-null -e hindIII --adapter-1 ACACTCTTTCCCTACACGACGCTCTTCCGATCT --adapter-mm 2 --len-limit 90)*. Overall, 85.3% reads passed all quality filtering steps and were deemed high-quality **(File S1)**. These reads were mapped to the *S. habrochaites* LYC4 genome (Aflitos *et al*., 2014) using BWA MEM v0.7.17 (Li, 2013) with default parameters. Resulting SAM files were converted to BAM and sorted using Samtools v1.9 (Li *et al*., 2009). Variant calling was performed with Stacks v2.3b using the default parameters for the reference-based mapping pipeline. Unfiltered SNPs were exported using the *populations* module with default parameters. Filtering of SNPs was performed with vcftools v0.1.15 (Danecek *et al*., 2011) using the following parameters *(--max-missing 0*.*8 --min-meanDP 6 --max-meanDP 30 --maf 0*.*05 --mac 3)*. All individuals had less than 50% missing loci, so none were removed. Heterozygosity and mean read depth were calculated for each sample in vcftools, which resulted in sample LA0716 being removed from downstream analyses due to higher than expected levels of heterozygosity. To make some downstream analyses easier to complete, linkage disequilibrium filtering was performed using plink v1.90b3.38 using 10 kb sliding windows and a r^2^ of 0.2 following a LD decay plot generated in PopLDdecay (Zhang *et al*., 2019). Two VCF files, the filtered only and filtered with LD pruning, were imported back into Stacks to produce the necessary input files for downstream analyses and calculating F statistics (F_st_). Accession-wise details are provided in **File S1**.

### Inference of ancestral population number

Population structure was assessed with two different approaches — using inference of ancestral populations and using coalescent analyses. Ancestral population estimation was performed using three different datasets for increased robustness: **(Set 1)** We assessed 254,263 SNPs using the R package LEA v2.4.0 (Frichot & François, 2015). STRUCTURE (Pritchard *et al*., 2000) and ADMIXTURE (Alexander *et al*., 2009) rely on simplified population genetic hypotheses such as the absence of genetic drift, as well as Hardy-Weinberg and linkage equilibrium in ancestral populations. LEA does not rely on the same assumptions and is more appropriate for inbred lineages (Frichot *et al*., 2014) and was therefore used due to the high-levels of self-compatibility found in some populations of *S. habrochaites*. **(Set 2)** Set 1 was further filtered using LD pruning as described above to produce a total of 93,129 SNPs, and analyzed using LEA. **(Set 3)** A different run of Stacks was performed using Stacks v1.44, which allowed more granularity in parameter selection. The non-default parameters included *(-T 3 -m 5 -S -- bound_low 0 --bound_high 0*.*02 --alpha 0*.*05)*. The *populations* module in Stacks v1.44 was called with the following non-default parameters *(-t 3 -r 0*.*5 -m 5 --min_maf 0*.*1 --lnl_lim -6 --merge_sites –write_random_snp*). This set contained 25,752 SNPs. For ancestral populations inference, analyses of K=2-15 were performed to determine the best K using the cross-entropy criterion (Frichot *et al*., 2014), using an alpha value of 100, and 200 iterations. Principal Component Analysis (PCA) as implemented in SNPrelate v1.16.0 (Zheng *et al*., 2012) was performed using Set 1 and 2 SNPs. Two PCAs were performed for each set: the full data set with all individuals included and one with the *S. pennellii* outgroups (LA1674, LA1809, and LA1941) removed.

### Inference of population relatedness using coalescent analysis

Coalescent analyses were performed with SNAPP as implemented in BEAST2 v2.4.5 (Bouckaert *et al*., 2014). Due to the computationally intense nature of SNAPP, the pruned SNP set was further pruned using vcftools by only including sites with no missing data and thinning SNPs to have a minimum of 50,000 bp between successive SNPs. This kept 3,965 SNPs. The resulting VCF file was then converted to a fasta file using vcf2phylip (Ortiz, 2019). The XML file was created using BEAUTi keeping each individual as unique species/populations. The mutation rate U and V were calculated from the data set with a coalescence rate set to 10. The MCMC was run for 8 million generations to achieve suitable ESS values (∼449). Tree visualization was performed with DensiTree (part of the Beast package) using a 25% burnin. The maximum clade credibility (MCC) was also identified with TreeAnnotator (part of the Beast package) using a 25% burnin. The tree allowed for classification of eight clades within the ingroup (plus the outgroup). These clades were then used for downstream analyses that required population specification. The MCC was then plotted on a map along with heterozygosity levels using the R package phytools v0.6.99 (Revell, 2012).

### Other population genomic analyses

Isolation by Distance analyses was performed using Mantel’s test of correlation between the genetic and geographic distance matrices using Set 2 SNPs. The genepop file produced by Stacks was read into the R package adegenet v2.1.1 (Jombart, 2008; Jombart & Ahmed, 2011) as each accession being a unique population. Two sets of analyses were done: one with all the accessions found in the coalescent analysis (minus the outgroup) and a second set of analyses with samples split between northern and southern super-clusters as seen with the MCC plotted on a map. Both analyses used Euclidean distances and 100,000 permutations. To investigate historical migration patterns, TreeMix v1.13 (Pickrell & Pritchard, 2012) was used. Samples were separated into the eight categories identified by SNAPP plus the outgroup. Overall, zero to ten migration events were tested with a likelihood ratio test done to determine which migration events were significant **(Table S1)**. Variance explained upon no migration and addition of individual migration events was obtained using the R function *get_f* in Treemix.

### Identification of SNPs for Targeted Sanger Sequencing

Analysis of reproductive traits identified populations of significant phenotypic interest that were not included in the original population genetic analysis. Thus, 15 accessions not included in the RAD-seq study (as well as three control accessions in the RAD-seq study) **(Table 1)** were analyzed using targeted Sanger sequencing (TSS) of 22 polymorphic loci, which were selected as follows: Since Stacks v2.x does not provide information about the breadth and coverage of individual SNPs, we used Stacks v1.44 to obtain SNP catalogs using **Set 3 SNPs** as described above. Custom Python scripts were used to identify 36 high-confidence polymorphic loci that were present across at least 50 out of 51 accessions, had a read coverage of >10X, and were not blacklisted by the *populations* module for STRUCTURE analysis. The broad coverage across almost all accessions was intended to ensure most of them would be captured in any novel set of accessions using targeted sequencing. Twenty four of these 36 loci were randomly selected and using their genomic locations and 100 base regions on either side of the 100 base RAD-tag were extracted for primer design. Amplicons could be successfully obtained for 22 loci.

**Table 1:** Reproductive traits for *S. habrochaites* accessions. Reproductive traits documented for each accession include mating system as detected by fruit production after self-pollination and/or pollen tube growth analysis (SC = self-compatible, SI = self-incompatible); expression of S-RNase protein as detected by immunoblotting; *S-RNase* allele as detected by allele-specific PCR with at least three individuals in each accession; presence of HT protein as detected by immunoblotting; pollen-side interpopulation reproductive barriers as detected by pollen tube growth in crosses with pollen from different accessions onto pistils of SI accession LA1777 (Y = pollen tubes rejected, N = pollen tubes accepted); pistil-side interpopulation barriers as detected by pollen tube growth with pollen of SC accession LA0407 onto pistils of different accessions (Y = pollen rejected, N = pollen accepted); inter-specific unilateral incompatibility (UI) detected by pollen tube growth in crosses using cultivar (*S. lycopersicum*) pollen onto pistils of different accessions (Y = pollen tubes rejected, N = pollen tubes accepted); and SC type based on the combination of reproductive traits and S-RNase allele present. *Data from Broz et al., 2017, ^Data from Covey et al., 2010, ^#^Data from Markova et al., 2016, ^a^presence of *LhgSRN-1* allele detected at a low frequency, nt = not tested, na = not applicable because accession is SI. Unshaded portion of the Table shows accessions from Ecuador and shaded portion of the Table shows accessions from Peru.

| Accession | Mating System | S-RNase protein | *S-RNase* Allele(s) | HT protein | Inter-pop pollen-side | Inter-pop pistil-side | UI | SC type |
| --- | --- | --- | --- | --- | --- | --- | --- | --- |
| LA4656/<br>ECU1498 | SC | N | *LhgSRN-1* | Y | Y | N | Y | 2 |
| LA1624* | SC | N | *LhgSRN-1* | Y | Y | N | Y | 2 |
| PI129157* | SC | N | *LhgSRN-1* | Y | Y | N | Y | 2 |
| LA1625* | SC | N | *LhgSRN-1* | Y | Y | N | Y | 2 |
| LA1266* | SC | N | *hab-7* | Y | N | N | Y | 1 |
| PI134417 | SC | N | *LhgSRN-1* | Y | Y<sup>#</sup> | N | nt | 2 |
| LA1264* | SC | N | *hab-7* | Y | N | N | Y | 1 |
| PI390515 | SC | N | *LhgSRN-1* | N | Y | N | N | 3 |
| LA0407* | SC | N | *LhgSRN-1* | Y | Y | N | Y | 2 |
| LA1223* | SC | N | *LhgSRN-1* | N | N | N | N | 3 |
| PI251305 | SC | Y | *hab-7*/<br>*LhgSRN-1* | N | N | Y | Y | 1 |
| LA4654/<br>ECU434 | SC | N | Unknown | Y | N | N | Y | 6 |
| LA4655/<br>ECU436 | SC | N | Unknown | N | N | N | Y | 6 |
| LA2119* | SC | N | *hab-7* | Y | N | N | Y | 1 |
| LA2868* | SI | Y | Multiple<sup>a</sup> | Y | N | Y | Y | na |
| LA2128 | SC | N | *hab-7* | Y | N | N | Y | 1 |
| LA1252 | SC | N | *hab-7* | Y | N | N | Y | 1 |
| LA2855 | MP | Y | Multiple | Y | N | Y | Y | na |
| LA2106* | SC | N | *hab-7* | Y | N | N | Y | 1 |
| LA2101* | SC | N | Unknown | Y | N | N | Y | 5 |
| LA2860 | SC | N | Unknown | Y | N | N | Y | 5 |
| LA2864* | SI | Y | Multiple | Y | N | Y | Y | na |
| LA2099* | MP | Y | Multiple<sup>a</sup> | Y | N | Y | Y | na |
| LA2098* | MP | Y | Multiple | Y | N | Y | Y | na |
| LA2175* | MP | Y | Multiple | Y | N | Y | Y | na |
| LA1391* | MP | Y | Multiple<sup>a</sup> | Y | N | Y | Y | na |
| LA2314* | SI | Y | Multiple | Y | N | Y | Y | na |
| LA1353*<sup>^</sup> | SI | Y | Multiple | Y | N | Y | Y | na |
| LA1777*<sup>^</sup> | SI | Y | Multiple | Y | N | Y | Y | na |
| LA0094 | MP | Y | Multiple<sup>a</sup><br>inc *hab-6* | Y | N | Y | Y | na |
| LA1691 | SC | Y | *hab-6* | N | N | nt | Y | 4 |
| LA1681 | SC | Y | *hab-6* | Y | N | nt | nt | 4 |
| LA1721 | SC | Y | *hab-6* | Y | N | Y | nt | 4 |
| LA1927<sup>^</sup> | SC | Y | *hab-6* | Y | Y | Y | Y | 4 |

### Targeted Sanger Sequencing for population structure

An initial nucleotide BLAST was performed using the *LYC4* (Aflitos *et al*., 2014) *S. habrochaites* assembly to identify sequences surrounding the polymorphic sequences for each of the 22 loci. Primers were designed to specifically amplify ∼200 bp spanning the polymorphic regions of each locus, and the resulting PCR products were purified (Zymo, Irvine,CA), and sequenced (GENEWIZ, South Plainfield, NJ). For each of the 22 loci, sequences were aligned in MEGA7 (Kumar *et al*., 2016) using Muscle (Edgar, 2004) with the original intended target, the corresponding sequence of LA0407 from the original RAD-seq dataset, and the top BLAST hits.

The diploid state of each locus of each accession was determined by aligning the sequences for all accessions, trimming off poor-quality sequences, and examining the set of trace files for each locus manually for heterozygous base calls (Chromas Pro, https://technelysium.com.au/wp/chromaspro/). If no ambiguous calls were present in the trace files, the individual was assumed to be homozygous at that locus.

To combine TSS and RAD-seq data, we performed BLAST between the trimmed Sanger sequences and the set of all 22 RAD loci consensus sequences to identify each corresponding RAD-seq locus. Sequences representing the 22 loci were extracted from the RAD-seq data (51 samples) in the *populations* module of STACKS v1.46 using a selection of loci identified by BLAST. These sequences were combined with their targeted Sanger sequencing counterparts using custom scripts, aligned, manually inspected, and trimmed when necessary. PGDSpider v2.1.1.2 (Lischer & Excoffier, 2012) was used to call allele variants for each locus; the separate matrices generated by PGDSpider were then combined to create the STRUCTURE v2.3.4 (Pritchard *et al*., 2000) input matrix of allele variants for all 22 loci for the 69 total samples (15 Sanger sequences from accessions not in the original experiment, 3 Sanger sequences from accessions included in the original RAD-seq dataset, and 51 sequences from the RAD-seq samples) using a custom Python script. STRUCTURE was run using default parameters with no prior population groups assumed for K=1-8 (three replicates per K) for 10,000 burn-in and 10,000 MCMC cycles. Results were extracted using STRUCTURE HARVESTER vA.2 July 2014 (Earl and von Holdt, 2011) and replicate runs were combined using CLUMPP v1.1 (Jakobsson & Rosenberg, 2007). All statistics (adegenet v1.7-15, pophelper v2.3.0), data analysis (pophelper), and plot generation (ggplot2, scatterpie) were performed using R v3.4.1 (Jombart & Ahmed, 2011; Francis, 2017).

### Identification of S-RNase allele hab-7

A previously published stylar transcriptome of SC accession LA2119 (Broz *et al*., 2017a) was used as a BLAST database to discover potential *S-RNase* alleles (NCBI BioProject PRJNA310635). Using a set of known *S-RNase* gene sequences as BLASTn queries to the LA2119 assembly, a single putative *S-RNase* transcript sequence was recovered. Allele-specific primers were designed using this putative *S-RNas*e sequence **(File S2)**, and PCR was performed using genomic DNA from multiple LA2119 individuals. Amplicon sequencing verified the sequence identified by the transcriptome analysis and revealed the presence of a single intron in genomic DNA. Following the convention set by Covey *et al*. (2010), this S-RNase allele was dubbed *hab-7*. The transcript abundances of the *hab-7* allele in LA2119 styles and two different *S-RNase* alleles of SI *S. habrochaites* accession LA1777 were identified using data from a previous *S. habrochaites* transcriptome study (Broz *et al*., 2017a).

### Reproductive trait analysis

At least two genetically distinct individuals (each grown from a separate seed) of each accession were used for phenotyping. Mating system was determined for previously untested northern accessions **(Fig. S5)** and verified in an additional set of accessions using self-pollinations as previously described (Broz *et al*., 2017b). If production of self-fruit was observed, plants were recorded as SC. If plants failed to set self-fruit using this approach, hand pollinations were performed, and/or pollen tube growth in styles was assessed as previously described (Covey *et al*., 2010). When at least three pollen tubes could be visualized at the base of the style or in the ovary in multiple independent crosses, plants were considered SC. When no self-fruit was formed and pollen tube tips could clearly be visualized terminating within the style, plants were considered SI. To test for inter-population reproductive barriers as initially described by Martin (1961), hand pollinations were performed using *S. habrochaites* SC accession LA0407 (SC-2 group) as male to test for the presence of pistil barriers that reject pollen of SC-2 plants and SI accession LA1777 as female to test for pollen resistance to S-RNase barriers as previously described (Broz *et al*., 2017b). To test for interspecific reproductive barriers, pistils of *S. habrochaites* accessions were pollinated using *S. lycopersicum* cultivars VF36, M82 or LA1221 as males.

Expression of *S-RNase* and an additional pistil SI factor, HT-protein, was assessed in stylar extracts from at least 2 individuals using immunoblotting with anti-peptide antibodies specific to each protein as described previously (Covey *et al*., 2010; Chalivendra *et al*., 2013; Broz *et al*., 2017b). The presence or absence of specific *S-RNase* alleles was determined for at least three individuals from each accession. *S-RNase* alleles were amplified from genomic DNA of individual plants using allele-specific primers **(File S2)** in PCR reactions, as described previously (Broz *et al*., 2017b). In selected accessions, the *HT* gene was amplified from genomic DNA using conserved gene specific primers (Covey *et al*., 2010), PCR products were purified and subjected to Sanger sequencing.

### Acylsugar sampling and MS data analysis

Acylsugar sampling was performed from a single uniformly sized young leaflet of 2-3 plants per accession for 40 of the 50 accessions as previously described (Fan *et al*., 2016). Metabolites were analyzed on a Supelco Ascentis C18 column, using a Shimadzu Ultra High Performance Liquid Chromatograph (UHPLC) connected to a Waters Xevo quadrupole Time of Flight mass spectrometer (MS). Raw files were converted into ABF format using Reifycs Abf Converter (https://www.reifycs.com/AbfConverter/) and imported into MS-DIAL (Tsugawa *et al*., 2015) for preprocessing. Peaks with an amplitude >100 and with >2 data points were considered for further analysis. Mass slice width and sigma windows were set to 0.05 and 0.5, respectively. Peaks across all samples were aligned with a 0.05 min. retention time tolerance and 0.03 Daltons MS1 tolerance. Acylsugars were then selected and annotated from the alignment results based on manually identified acylsugar peaks, MS1 *m/z* values, MS/MS and previous results (Kim *et al*., 2012). A five-fold sample max/blank average filter was applied across the samples. Normalized and filtered data was then exported. Extracted peak areas were normalized by the internal standard peak areas per sample, and the normalized peak areas were averaged for each accession. Only peaks >2X internal standard peak area in >2 accessions were considered reliable signals.

## Results

### Preliminary analysis of RAD-seq data

For RAD-seq, we selected 52 out of the >100 accessions of *S. habrochaites* in the TGRC germplasm database (https://tgrc.ucdavis.edu/) that span the entire species range. Four *S. pennellii* accessions were included as outgroups. As expected, there was large variation in the number of reads obtained per sample **(Fig. S1A)**, ranging from 5707 to 5,348,246. After filtering the initial reads based on quality, one or both replicates of five accessions – LA2868, LA2976, LA1978, LA2098, LA0716 – were removed **(Fig. S1A)**, still leaving 53 accessions of the two species for further analyses **(File S1)**. We did not find any correspondence between number of reads between technical replicates **(Fig. S1B)** suggesting that the read number variation was due to the randomness associated with restriction cleavage and the RAD-seq library preparation protocol as opposed to within-species variation of restriction sites. To improve read coverage per accession, we combined reads from the technical replicates and mapped all reads to the draft-quality *S. habrochaites* LYC4 genome (Aflitos *et al*., 2014) to make clustered read stacks (Catchen *et al*., 2013). The number of retained loci and mean coverage across all loci also varied across accessions **(Fig. S1C**,**D)**. The median read coverage across all sites after filtering was 9.5, with 20 out of 53 accessions having a mean read coverage ≥10X **(Fig. S1D)**.

### Defining the population structure in Solanum habrochaites

To assess the genetic relatedness between *S. habrochaites* accessions across the range, we first inferred SNP-genotype based ancestral populations (K) using three different SNP datasets. These results suggested that the sampled *S. habrochaites* individuals arose from 4-6 ancestral populations **(Fig. 1; Fig. S2)** – three of which (*purple, red, yello*w) lie south of the AHZ. In the Ecuador, the optimal K=4 suggested presence of one ancestral *orange* population **(Fig. 1A**,**B)** while K=6 discriminated individuals in this region as originating from two different ancestral populations – *orange* and *green* – with different degrees of genotype sharing between them **(Fig. 1D)**. Manual observation of population assignments at multiple Ks **(Fig. S2)** as well as results of coalescent analysis **(Fig. 2A)**, targeted Sanger sequencing **(Fig. 4)** and previous results using short sequence repeat markers (Sifres *et al*., 2011), led us to assign six ancestral populations **(Fig. 1D)**. We also observed that multiple accessions – at both K=4 and K=6 values – across the range had small yet non-negligible probabilities of being assigned to *yellow, red* and *purple* populations, south of the AHZ.

**Figure 1:**
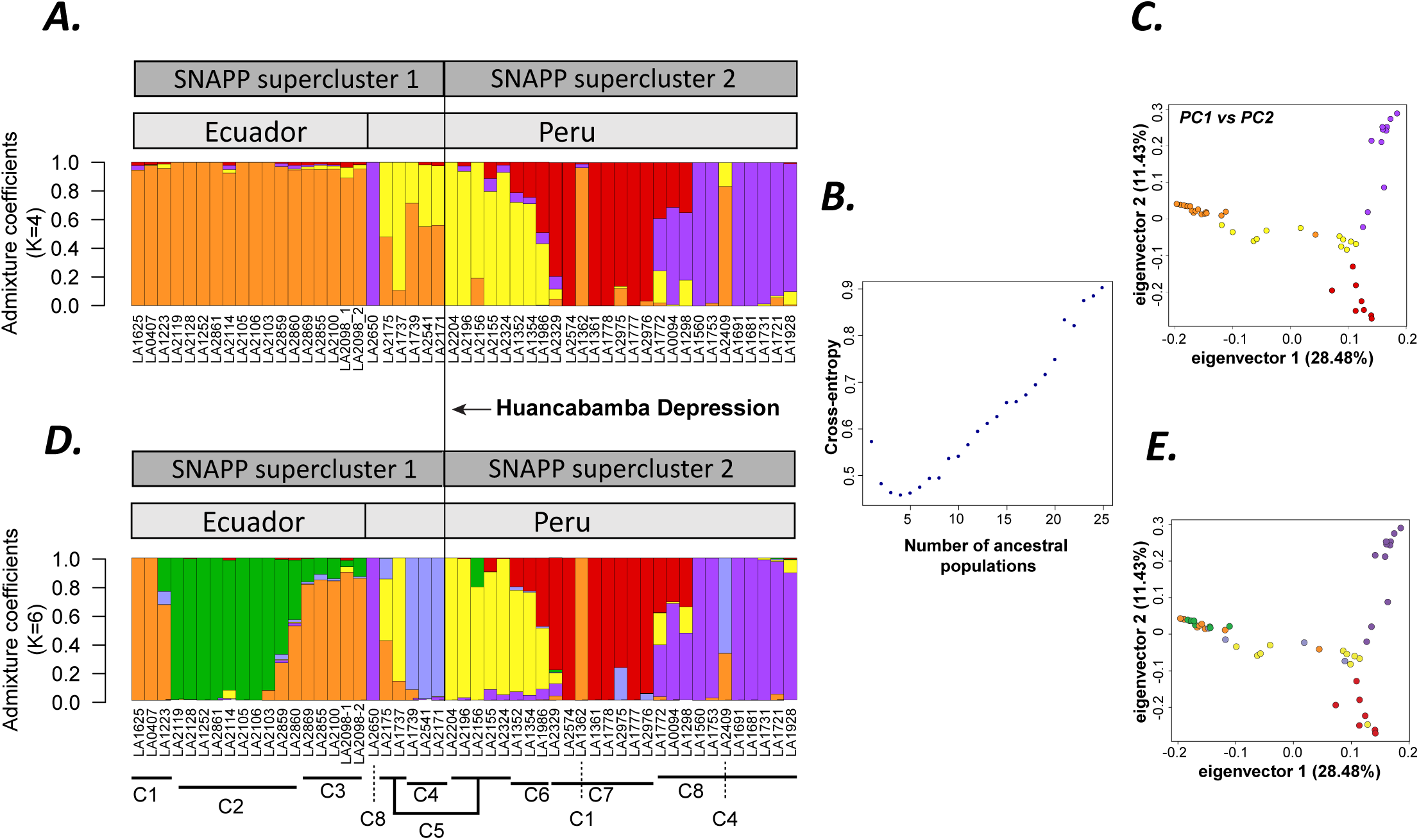
Population structure of S. habrochaites. (A, D) Population structure plots obtained using K=4 and K=6 as pre-defined population clusters using Set 1 SNPs. Population cluster numbers as per Fig. 2A are described below the barplot in (D). (B) Cross-entropy criterion showing K=4 as the optimal number of genetically differentiated ancestral populations. (C, E) Principal Components Analysis, with populations defined based on K=4 and K=6 as shown in sub-figures A,D, respectively.

PCA using Set 1 and 2 markers **(Fig. 1C**,**F; Fig. S3)** showed a close relationship between accessions north of the HD, but interestingly in the south, the southernmost *purple* individuals were more related to *yellow* individuals near the central part of the range – near Huancabamba/Cajamarca – than to their geographically close *red* individuals. To better understand this observation, we employed a coalescent tree-based approach using SNAPP (Bryant *et al*., 2012), with 3965 markers represented in all *S. habrochaites* and *S. pennellii* accessions. This algorithm infers Bayesian trees for every SNP identified in the population and integrates for coalescence across all trees. Combined trees across all SNPs identified eight genotypically distinct population clusters within *S. habrochaites* **(Fig. 2A)**, largely following the populations identified by LEA at K=6 **(Fig. 1D)**. The two additional clusters 3 and 6 comprised of genotypic hybrids between the *green-orange* and *red*-*yellow* populations **(Fig. 1D)**. Population clusters 1-4 (supercluster 1; northern supercluster) correspond to samples from mid/southern Ecuador and northern Peru, while clusters 5-8 (supercluster 2; southern supercluster) correspond to samples across Peru **(Fig. 1F)**. The two superclusters are separated at the HD, suggesting the geographical feature’s role in modulating species dispersal. Unexpectedly, we also found that cluster 8 was not closely related to its geographical neighbor cluster 7 but was genetically equidistant from clusters 5-7.

**Figure 2:**
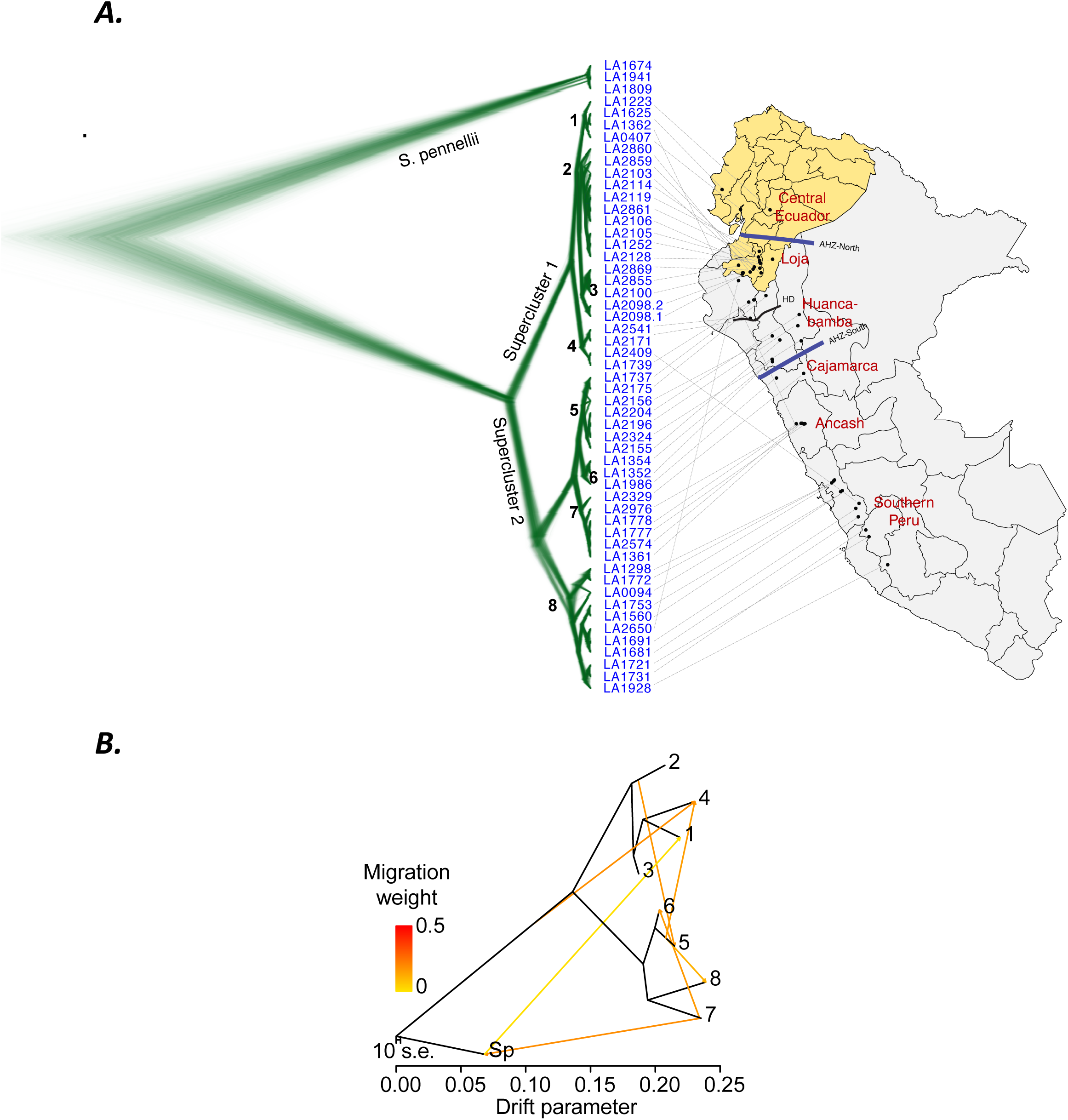
Coalescent and migration analysis. (A) Results of coalescent analysis using SNAPP, obtained using markers shared between all sampled individuals. LA2975 was left out from this analysis because its level of heterozygosity was >3X the next highest sample, suggesting possible contamination or other unexplained behavior. AHZ: Amotape-Huancabamba Zone. HD: Huancabamba Depression. Population cluster numbers are marked within the phylogeny. (B) TreeMix analysis showing inferred migration events between different clusters. Migration weight indicates confidence in a given inferred migration event. Tree obtained in (A) is using coalescent Bayesian analysis while that in (B) is using maximum likelihood analysis by the individual software packages.

Taken together, the three methods suggest that there are eight current population clusters across the sampled *S. habrochaites* accessions, likely derived from 4-6 ancestral populations. Three clusters (clusters 6-8) remained robust to the different analyses performed, indicating that they are more genetically distinct. Cluster 8 – which was largely SC except for LA1298 (SI), LA1772 (SI), LA0094 (MP/SI) (see mating system analysis below) – was genetically equidistant and well-separated from clusters 5-7. Unexpectedly, the three SI accessions bore evidence of a shared genotype with the *yellow* and *red* populations. We explored three scenarios to explain this set of observations better. First, we speculated that the three accessions could be a result of hybridization between a SI *red*/*yellow* genotype male and a SC *purple* genotype female. Analysis of migration between populations using a phylogenetic co-variance-based approach (Pickrell & Pritchard, 2012) suggested presence of eight migration events (seven vs six events: D-statistic 17.78, p-value=2.49e-05; eight vs seven events: D-statistic -3.76, p-value=1) **(Fig. 2B; Table S1)**, of which one event is between clusters 5 and 8. However, evidence of such a migration would leave a significant footprint in the coalescent tree, which is not observed. Second, we asked whether these three individuals represent an ancestral SI population from which the *yellow, red* and *purple* populations evolved. This scenario would result in clusters 5,6 being more closely related to clusters 3,4, which is not the case. Also, previous studies suggest *S. habrochaites* origin to be further north, near the AHZ (Rick *et al*., 1979; Sifres *et al*., 2011; Pease *et al*., 2016). Thus, we explored a third scenario **(Fig. 6)**. Here, cluster 8 would have been established by southward migration from near the AHZ (clusters 5,6), from founders possessing *red, yellow* and *purple* genotypes. Under this scenario, cluster 7 would result from an independent, local fixation of the *red* genotype. This hypothesis is supported by PCA, which suggests a closer relationship between cluster 8 and clusters 5,6 than with cluster 7 **(Fig. S3)** as well as by presence of small *red, yellow* and *purple* genotype probabilities in Ecuadorian accessions **(Fig. 1A**,**D)**. Further analysis with fixation index also supported this model (see population differentiation section below). This scenario also allows for a rapid separation of the two superclusters via northward/southward migrations, as seen in the coalescent analysis **(Fig. 2A)**. The third scenario is thus the most plausible, and also supports previous inferences of the origin of *S*.*habrochaites* after separation from *S. pennellii* near the Cajamarca/Huancabamba region **(Fig. S3)** (Rick *et al*., 1979; Sifres *et al*., 2011; Pease *et al*., 2016). By studying acylsugar phenotypes below, we further define the origin as the Cajamarca region at the southern border of the AHZ.

After *S. habrochaites*-*S. pennellii* divergence, some *S. habrochaites* individuals migrated northwards through the river valleys of the AHZ. The drastic separation between the two superclusters at the HD **(Fig. 2A)** suggests that there has been little gene flow between them. We thus sought to assess the impact of HD and other barriers to gene flow across the species range.

### Characterizing population differentiation in S. habrochaites

Three metrics were estimated – heterozygosity, fixation index (Fst) and gene flow due to migration events. Overall, heterozygosity levels in *S. habrochaites* ranged from 0.02-0.17, while that in *S. pennellii* ranged from 0.02-0.07 **(Fig. S4)**. Values for SI *S. pennellii* were lower than expected, possibly because two of the three accessions (LA1809 and LA1941) were collected at the extreme northern and southern species margins, respectively. As expected, *S. habrochaites* SC accessions had lower heterozygosity than SI accessions **(Fig. S4)**. The observed heterozygosity across the range was substantially lower than the expected heterozygosity (**Fig. S4)**, suggesting significant deviation from Hardy-Weinberg equilibrium, even in SI populations. This may be expected given many SI/MP accessions lie in the AHZ. A previous study (Sifres *et al*., 2011) concluded based on SSR markers that the highest heterozygosity existed among accessions from the Huancabamba and Cajamarca regions. We utilized the ecogeographic groups defined in that study to directly compare their heterozygosity with those from our RAD-seq data. Both the observed and expected heterozygosity values per accession were lower using RAD-seq than SSRs **(Fig. 3A)**, a trend that has been observed previously (Sunde *et al*., 2020). This may be due to the greater probability of identifying conserved restriction site linked markers in genome-wide RAD-seq studies as opposed to meaningful biological differences. Thus, the relative differences between populations are likely to be more important than specific values (Sunde *et al*., 2020). Indeed, the trend observed between ecogeographic groups here was similar to the previous SSR study (Sifres *et al*., 2011). The patterns of heterozygosity also correlated well with the mating systems – all Huancabamba, Cajamarca and Ancash accessions were SI/MP, and display higher levels of heterozygosity while the population extremes were SC with lower heterozygosity.

**Figure 3:**
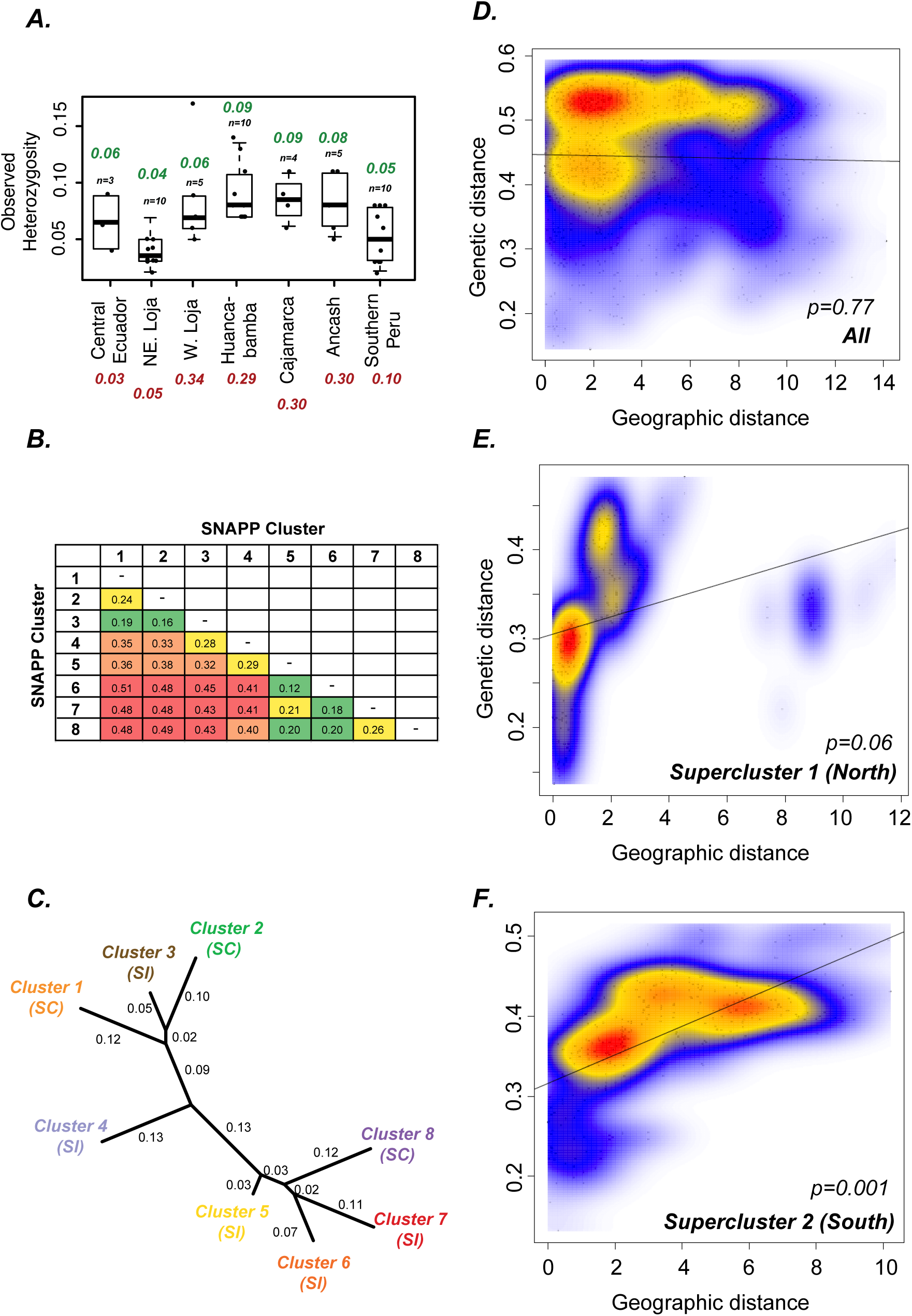
Analysis of population relatedness and demographic events. (A) Observed heterozygosity estimates for individuals classified by their geographic regions. Northeastern Loja and Western Loja all comprise individuals assigned to clusters 2 and 3, respectively. Number in green above the boxplot corresponds to the average heterozygosity from this study, while those in red below the region names correspond to SSR marker estimates of observed heterozygosity as per Sifres et al, 2011. Outlier value of LA2098-2 was excluded when calculating average for W. Loja. (B) Estimates of pairwise Fst between SNAPP coalescent clusters. Cells are colored as green (high Fst; <0.20), yellow (intermediate high Fst; 0.21-0.30); orange (intermediate low Fst; 0.31-0.40) and red (low Fst; >0.40). (C) Dendrogram based on the Fst matrix shows differentiation between clusters that follows the two SNAPP superclusters. Branch-wise Fst values are shown (D-F) Isolation by Distance analysis considering all *S. habrochaites* individuals, as well as those in Supercluster 1 and Supercluster 2. Mantel’s test p-values were estimated using 100,000 simulated permutations of the LD pruned Set SNPs.

Fst quantifies the degree of genetic variance between two populations that can be attributed to population structure, with values close to 0 indicating frequent inter-breeding and higher values indicating differentiation (Wright, 1931; Weir, 2012). Fst values ranged from 0.12 (clusters 5-6) to 0.51 (clusters 1-6) **(Fig. 3B)**. Values between clusters 5/6-8 were substantially higher than that between clusters 7-8, supporting the model of the former clusters’ relatedness as described above. Nevertheless, all Fst values were high enough to indicate barriers to unrestricted gene flow between populations. One of these barriers is the AHZ/HD, given the differentiation seen between the two superclusters at the HD and the low Fst of cluster 4, which lies in the heart of AHZ. In addition, when the Fst values were organized into a cladogram, the northern and southern superclusters again separate at the HD **(Fig. 3C)**. Another barrier to gene flow is evolution of SC at the range extremes – the Fst cladogram **(Fig. 3C)** illustrates that the SC populations generally have higher branch lengths than SI populations they are closely related with. We also found that cluster 2, despite being SC, has substantial allelic similarity with SI cluster 3. In conjunction with results from coalescent and population structure analyses **(Figs. 1**,**2)**, this pattern suggests that the SC cluster 2 may have emerged from the SI cluster 3.

The high Fst branch length of SI cluster 7 also led us to hypothesize that geographical distance acts as another barrier to species dispersal and connectivity, accounting for population differentiation in the southern part of the range. Isolation by Distance (IBD) analysis across the entire range did not show a significant association between genetic and geographic distance when all the samples were included, with the observed association being -0.02 (Mantel’s test p=0.77) **(Fig. 3D)**. When northern and southern superclusters were analyzed separately, the northern accessions showed a slight positive yet non-significant association of 0.32 (Mantel’s test p=0.06) **(Fig. 3E)**, but the southern accessions – all south of the HD – showed a significant IBD with an observed association of 0.63 (Mantel’s test p=0.001) **(Fig. 3F)**, supporting our hypothesis. Geographical distance, thus, is another barrier to gene flow between populations south of the HD. Combined together, these results reveal the role of three important players in *S. habrochaites* population differentiation – the evolution of SC, AHZ/HD, and the large geographical distances south of the HD. We identified different migration events **(Fig. 2B)**, but 99.1% of the genetic variance between populations could be explained by phylogeny alone **(Table S1)**, suggesting migration did not play a major role in population differentiation.

Cluster 4, close to the HD, was found to be unique, given that it is experiencing significantly restricted gene flow despite being SI. This pattern could be an outlier, being based on only three *bona fide* Huancabamba accessions **(Fig. 2A)**, however, given the high endemism in the AHZ, it could also be a result of geographic barriers to dispersal. Genome-wide differentiation is also clearly seen in the SC populations. There is evidence of genotype sharing in some accessions mostly in the contact zones between the SC/SI clusters, likely as a result of shared ancestry **(Fig. 1A**,**D; Fig. 3B)**. These results show that emergence of SC has clearly contributed to evolution of *S. habrochaites* diversity. We thus sought to characterize the mechanisms behind emergence of SC in this species..

### Reproductive traits in S. habrochaites accessions

*S. habrochaites* displays substantial diversity in the expression of reproductive traits including mating system, reproductive barriers between *S. habrochaites* populations and reproductive barriers between *S. habrochaites* and other tomato clade species. We combined new phenotyping results for ten Ecuadorian accessions **(Fig. S5)** with data from previous studies (Martin, 1961, 1963; Rick *et al*., 1979; Mutschler & Liedl, 1994; Sacks & St. Clair, 1998; Covey *et al*., 2010; Baek *et al*., 2015; Markova *et al*., 2016; Broz *et al*., 2017b) (https://tgrc.ucdavis.edu/) to generate a comprehensive inventory of mating system and other reproductive traits throughout the *S. habrochaites* range **(Table 1)**.

An SC mating system was confirmed for 19 accessions at the northern species margin in Ecuador **(Table 1)**. In general, our results were congruent with previous reports (Rick *et al*., 1979) or with mating systems designated by the TGRC (https://tgrc.ucdavis.edu/), with a few exceptions. The Ecuadorian accession LA2855, which had previously been designated as facultative SC, was found to contain both SI and SC individuals (now designated mixed population; MP). At the southern species range margin, accession LA0094 was also found to be MP (previously designated as SC), and accessions LA1753 and LA1560 were found to be SC rather than SI. In addition, we determined that all tested accessions in cluster 3 are MP, while those tested in cluster 2 are SC.

The pistil barrier/pollen resistance architecture of S-RNase-based gametophytic SI suggests that typically, SI will be lost due to mutations in pistil-expressed genes, e.g. due to loss of *S-RNase* and/or loss of *HT* expression in styles (Bedinger *et al*., 2017). We assessed the expression of S-RNase and HT proteins in stylar extracts using immunoblotting (**Table 1; Fig. S5B**). In general, S-RNase proteins were not detectable in styles of the northern SC accessions, congruent with previous reports (Broz *et al*., 2017b). Surprisingly, one northern SC accession (PI251305) expressed S-RNase(s) in three of four individuals tested. All SI and MP accessions expressed S-RNases, as expected. The southern marginal SC accessions that were tested do express S-RNase protein, consistent with previous studies that identified a low-activity S-RNase (hab-6) in southern SC accession LA1927 (Covey *et al*., 2010).

In several cases, SC could be correlated with specific *S-RNase* alleles **(Table 1)**. Previously, three distinct SC groups (designated SC-1, SC-2 and SC-3) were identified at the northern range margin based on inter-population and interspecies crossing behavior (Broz *et al*., 2017b); two of which (SC-2 and SC-3) were associated with the presence of S-RNase allele *LhgSRN-1*, which is not expressed at the RNA or protein level (Kondo *et al*., 2002; Covey *et al*., 2010; Broz *et al*., 2017b). The *LhgSRN-1* allele was detected in nine northern SC accessions in western coastal to central mountainous regions of Ecuador **(Table 1)**. The *LhgSRN-1* allele was also detected at a low frequency in several SI/MP accessions across the species range (LA2868, LA2099, LA1391, LA1353 and LA0094), suggesting that this “selfing allele” existed in the last common ancestral population of *S. habrochaites*, and was perhaps converted to an inactivated, non-expressed form in the AHZ during the species’ northward migration, before becoming fixed in the northern SC populations.

In this study, a new *S-RNase* allele (*hab-7*) was identified using stylar transcriptomic data from the SC-1 group accession LA2119 (Broz *et al*., 2017b,a). The *hab-7* allele **(Fig. S6)** appears to be a bona fide *S-RNase* allele, since it contains conserved features of all S-RNases including a predicted single intron, and a deduced amino acid sequence that harbors a secretory signal peptide, two catalytic histidine residues and the conserved C1-C5 motifs. In addition, the *hab-7* allele lacks sequences associated with non-S-locus S-like RNases (Vieira *et al*., 2008). Further, the hab-7 protein shares >99% identity with the available coding region of *S. peruvianum* S13-RNase (Chung *et al*., 1994) (GenBank: BAA04147.1). Expression of the *hab-7* allele in styles of SC accession LA2119 was 400-1200-fold lower than that of two different *S-RNase* alleles in styles of SI accession LA1777 (FPKM = 12.25 for *hab-7* in LA2119 versus 4604.90 and 14579.88 for the two S-RNases in LA1777). We found the *hab-7* allele in all known SC-1 group accessions and six geographically close accessions **(Table 1, Fig. 5)**.

A low activity S-RNase hab-6 was previously identified in southern SC accession LA1927 (Covey *et al*., 2010). We found the *hab-6 S-RNase* allele in all SC accessions tested at the southern *S. habrochaites* species margin (designated as SC-4).

Expression of HT protein – a pistil-specific factor not encoded at the *S* locus that functions both in SI and in interspecific pollen tube rejection (McClure *et al*., 1999; Tovar-Méndez *et al*., 2014; Tovar-Méndez *et al*., 2017) – was also assessed by immunoblotting. HT protein is expressed in styles of all tested accessions except for three from central Ecuador (LA1223, PI390515 and PI251305) and one from southern Peru (LA1691). PCR amplification and sequencing of the HT-A gene demonstrated that the three Ecuadorian accessions all have the same nonsense mutation in the second exon of the gene **(Fig. S7)**.

Unidirectional inter-population barriers between the northernmost SC and southernmost SC populations and between the extreme marginal SC and central SI populations have been documented in *S. habrochaites* (Martin, 1963; Rick & Chetelat, 1991; Markova *et al*., 2016; Broz *et al*., 2017b). We tested for pollen-side inter-population reproductive barriers by imaging growth of pollen tubes from different accessions in pistils of a well-characterized SI accession (LA1777) and found that, among our newly tested SC accessions, only LA4656 displays a pollen-side population barrier and thus exhibits all reproductive traits associated with the SC-2 group **(Fig. S5, Table 1)**. To assess pistil-side inter-population barriers, we tested growth of pollen tubes of a well-characterized SC-2 accession (LA0407) in pistils of the new accessions. We found that all accessions (either SI or SC) that express any S-RNase protein rejected LA0407 pollen and thus possessed pistil-side inter-population barriers (Broz *et al*., 2017b) **(Fig. S5, Table 1)**.

Finally, the presence or absence of interspecific reproductive barriers was assessed by examining the growth of interspecific pollen tubes in styles **(Fig. S5B)**. We tested for this trait using pollen from the cultivated tomato species *S. lycopersicum*, which is rejected in styles of wild tomato species that produce green fruits, including *S. habrochaites* (Baek *et al*., 2015). Most *S. habrochaites* accessions possess interspecific reproductive barriers, i.e. their styles reject pollen tubes of *S. lycopersicum*, but a previous study found a single accession (LA1223) that lacked this ability (Broz *et al*., 2017b). This accession also lacked expression of HT protein, did not reject pollen of SC-2 accessions and contained the *LhgSRN-1* S-RNase allele. This combination of traits was denoted as SC-3 (Broz *et al*., 2017b). In this study, we found only one additional SC accession, PI390515, which exhibited all of these traits and can be classified as a SC-3 group member **(Table 1, Fig. S5)**.

### Population identification of newly sampled accessions

Analysis of reproductive traits identified accessions of significant phenotypic interest that were not included in the RAD-seq population genetic study. To identify their ancestral populations, we utilized Targeted Sanger Sequencing (TSS) of 22 broadly represented and high coverage RAD-seq polymorphic loci. When the TSS and RAD-seq data for the 22 loci were combined and analyzed, seven ancestral populations could be identified that corresponded closely with reproductive characters (**Fig. 4A)**. One cluster identified using combined RAD-seq and TSS data (*orange*) corresponds to the coalescent cluster 1 **(Fig. 2A)** and includes accessions found along a steep altitudinal cline in central Ecuador **(Fig. 4)**. These accessions contain, or segregate for, the *Lhg-SRN1* allele found in the SC-2 and SC-3 groups **(Table 1)**. Notably, the northernmost SI accession in Ecuador, LA2868, which contains the *LhgSRN-1 S-RNase* allele at a low frequency **(Table 1)** also clusters with this group, and may represent an ancestral SI population. Another distinct cluster identified (*green;* **Fig. 4A**) includes SC accessions found in mountainous south-central Ecuador centered around Loja **(Fig 4B)**. These accessions all contain the low-expression *hab-7 S-RNase* allele. The accessions in this cluster that have been fully phenotyped for reproductive traits (Broz *et al*., 2017b) **(Table 1, Fig. S5)** lack inter-population reproductive barriers and exhibit interspecific barriers, consistent with a designation of SC-1 (Broz *et al*., 2017b) **Fig. S5)**. These SC-1 accessions all appear in coalescent cluster 2 in northeastern Loja **(Fig. S9)**.

**Figure 4.**
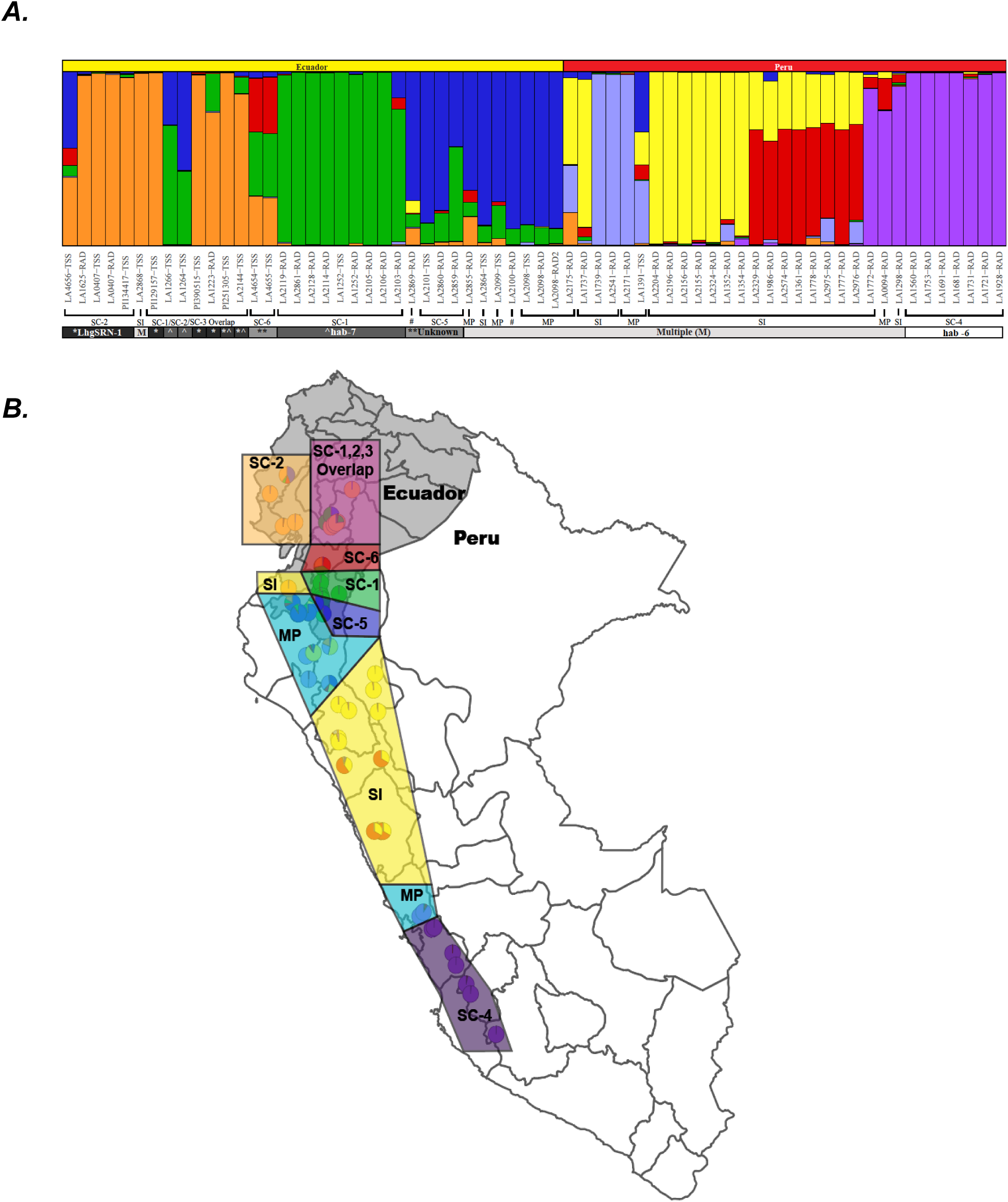
Mating system and population structure in Solanum habrochaites. (A) Population structure plot incorporating accessions analyzed using RAD-seq and targeted Sanger sequencing (TSS) organized in a north to south array from left to right. Mating systems indicated include SC groups 1-6 (Table 1), Mixed Population (MP) accessions containing both SI and SC individuals as well as purely SI accessions (SI). #, mating system not assessed. Where known, specific S-RNase alleles associated with accessions are indicated. *accessions containing the LhgSRN-1S-RNase allele, ^accessions containing the hab-7 S-RNase allele, **S-RNase allele unknown, and multiple (M) S-RNase alleles are found in SI and MP accessions. (B) Map of Ecuador and Peru displaying the locations of accessions with different mating systems as shown in (A) and listed in Table 1.

Unexpectedly, our *S-RNase* allele and *HT* expression data suggested that hybridization has occurred between the SC-1 and SC-2/3 groups, which overlap in a mountainous region near the town of Alausí **(Table 1, Fig. 4)**. Specifically, two accessions collected in this region (PI251305 and LA2144) include individuals containing either the *hab-7* or the *LhgSRN-1* alleles, or both. This result is corroborated by the full RAD-seq data for LA1223, an accession from the same region, where genetic relatedness to both northern and central Ecuador populations was seen using the filtered set of ∼95K markers **(Fig. 1D)**.

Two SC accessions collected west of the town of Gíron in central Ecuador LA4654 and LA4655 **(Fig. 4B)**, share a unique polymorphism pattern in the TSS analysis (**Fig. 4A**, *red/green/orange*). These accessions contain neither the *LghSRN-1* or the *hab-7 S-RNase* alleles, so *S-RNase* allele(s) are considered “Unknown” **(Fig. 4A, Table 1)**, and these accessions were tentatively designated as an “SC-6” group. These accessions lack inter-population barriers but retain interspecific barriers **(Table 1, Fig. S5)**.

Another cluster (*blue*, **Fig 4A**) contains SI and MP accessions as well as SC accessions with neither the *LghSRN-1* nor the *hab-7 S-RNase* allele (allele unknown, **Table 1**) and do not cluster with other SC types. Therefore, SC accessions in this cluster were tentatively designated as “SC-5” type. In the broader coalescent analysis with ∼95K markers, this cluster appeared as a hybrid between *orange* and *green* genotypes **(Fig. 1D)**, but is resolved into an independent population with 22 markers. The co-clustering of SC, MP and SI accessions in the *blue* cluster suggests that the emergence of distinct SC-5 and MP groups in southwest Ecuador region (**Fig 4B**) is recent enough that only moderate genetic differentiation has occurred between these groups and their SI progenitors.

In southern Peru, the SC-4 group with the *hab-6 S-RNase* allele (Covey *et al*., 2010) clusters with two SI accessions (LA1772 and LA1298) and one MP accession (LA0094) from the same region (**Fig. 4A**, *purple*). Given the finding of isolation by distance in supercluster 2, the emergence of the SC-4 group could have occurred from an MP type population (e.g. similar to LA0094) as a result of selection for reproductive assurance as the species migrated southward. This inference is supported by the coalescent analysis **(Fig. 2A)**, which shows LA0094 as more closely related to the SC accessions further south than other SI accessions in cluster 8.

### Evolution of acylsugar diversity in S. habrochaites

Results described above demonstrate the genetic differentiation between northern and southern population clusters brought about by multiple SI-->SC transitions and presence of the AHZ. Previous studies also identified differences in acylsugar profiles from different *S. habrochaites* accessions. One study (Kim *et al*., 2012) assessed the enzyme ASAT4 – the last enzyme in the acylsugar biosynthetic pathway – which was found to be inactivated in many northern accessions resulting in loss of acylsugar acetylation **(Figs. S8, S9)**. We re-analyzed this result in the context of population structure. As *S. habrochaites* sampled in this study were obtained from the stock center independently of the previous study, we re-sampled the leaf surface acylsugars from 2-3 replicates of 40/51 accessions used for RAD-seq. In two publications (Kim *et al*., 2012; Schilmiller *et al*., 2015), LA1362, LA2409 and LA2650 had been identified as chemotypic outliers in their geographical area i.e. their acylsugar phenotype matched not with their neighbors but with accessions very distant from their documented geographical area. We found that all three accessions were also genotypic outliers in their recorded geographic area **(Fig. 1, Fig. 2A)**. However, while LA2409 and LA2650 chemotypes matched previous results and their “true” geographic area based on their genotype, LA1362 profile did not appear as a chemotypic outlier in this study **(Fig. 5)**, the reasons for which are not currently clear.

**Figure 5:**
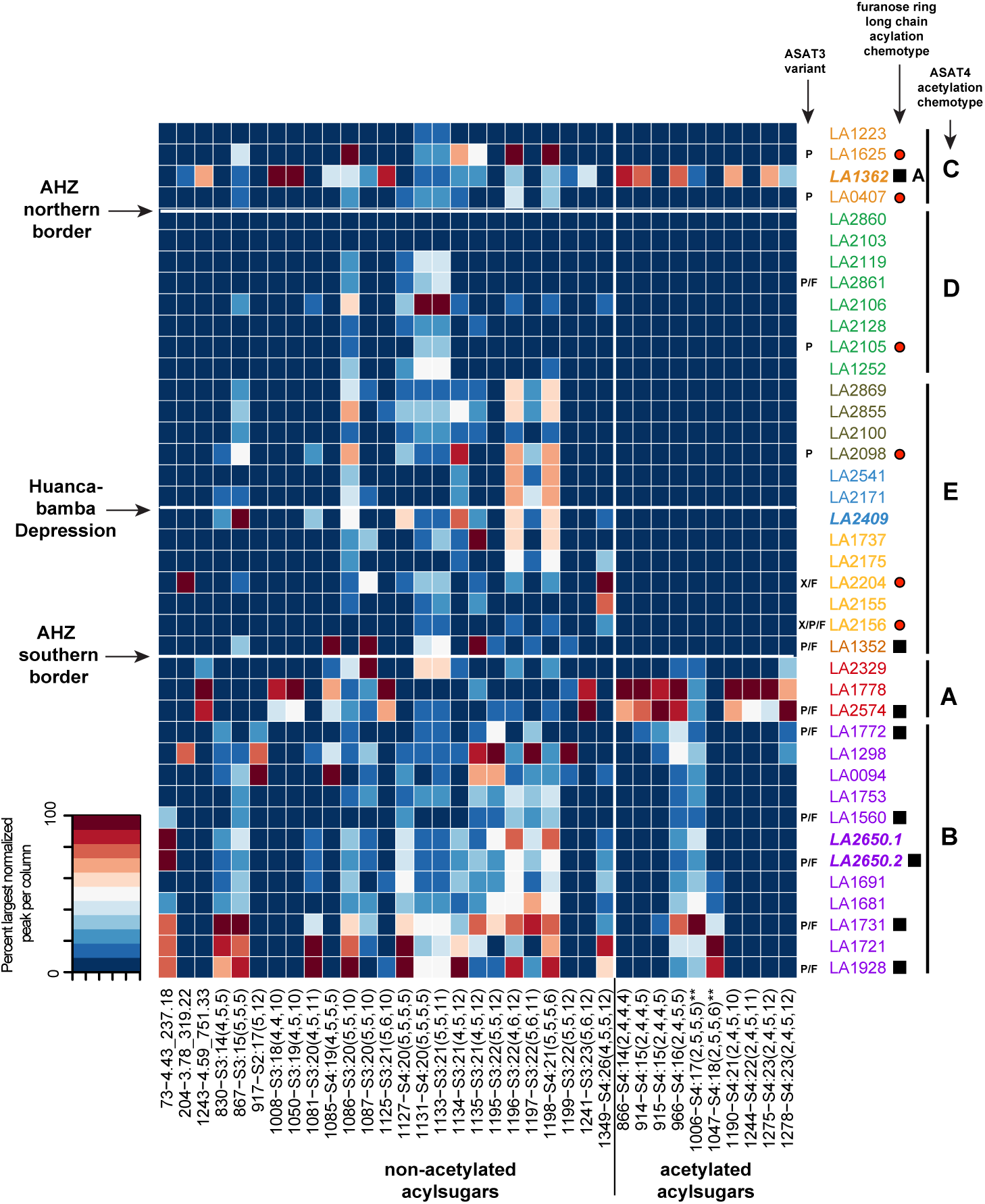
Acylsugar phenotypes across Solanum habrochaites accessions and the genotypes of two associated enzymes. Heatmap of acylsugar peak areas normalized to the internal standard peak area and the maximum area per column. Rows and columns were arranged based on Fig. 2A dendrogram and types of acylsugars, respectively. Accessions are colored by their population cluster assignments, using scheme used in Fig. 1D. Three accessions in bold are the geographically misplaced accessions.X/P/F indicate the three ASAT3 variants found in Schilmiller et al, 2015, while black squares and red circles indicate presence and absence of sucrose furanose ring long chain acylation as per the same study. Note that the black squares are associated with presence of both P/F while the red circles are largely associated with presence of only P. ASAT4 inactivation chemotypes (A,B,C,D,E) as per Kim et al, 2012 are also shown. Note that A,B have acetylated acylsugars and C,D,E contain only non-acetylated acylsugars, due to ASAT4 loss. Column names are in the format (peakID-identified acylsugar). Acylsugars with asterisks indicate those predicted based on MS1 peak and Kim et al, 2012 study without high-confidence MS/MS patterns.

Loss of acetylation in the north was previously found to occur via three different mechanisms – loss of *ASAT4* gene expression (chemotype E), frameshift mutation in *ASAT4* (chemotype D) and likely loss of the *ASAT4* gene (chemotype C) (Kim *et al*., 2012). Mapping these chemotypes onto the SI/SC definitions revealed that chemotypes C and D were confined to the northernmost SC clusters 1-6 (except SC-4, which is southern Peru), and chemotype E was widespread across SI/MP accessions abruptly limited by the southern boundary of the AHZ **(Fig. 5; Fig. S9)**. The two chemotypes A and B with acetylating, functional versions of ASAT4 – representing the ancestral ASAT4 activity – were associated with clusters 7 and 8. The northernmost accession with a functional *ASAT4* allele (LA2329) lies in the Cajamarca region south of the AHZ. This finding further supports the model of origination of *S. habrochaites* in the Cajamarca region, and thereby implies that the functional *ASAT4* allele was inactivated during the northward migration into the AHZ **(Fig. 6)**.

**Figure 6:**
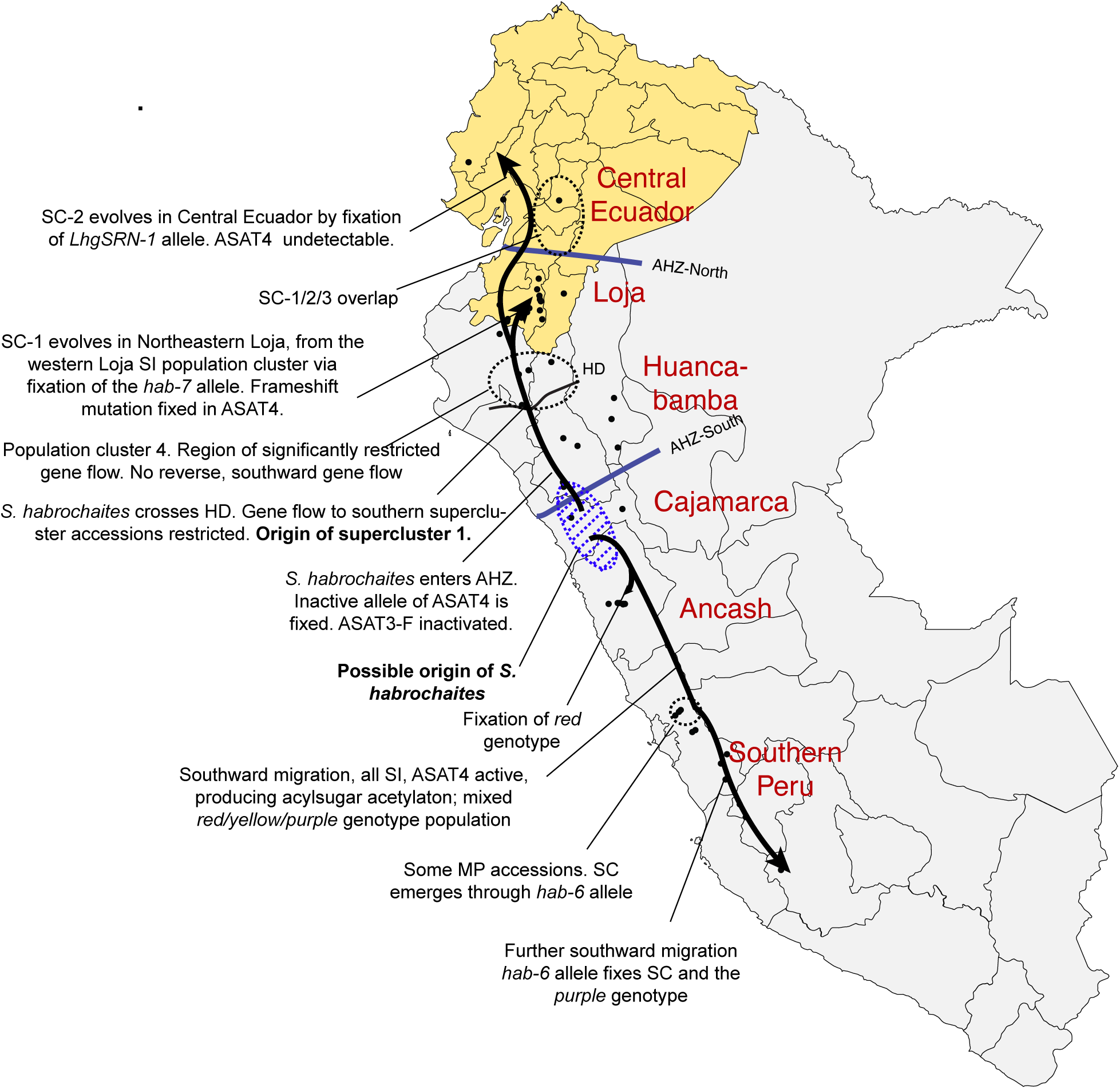
Overall model for Solanum habrochaites evolution. This model is based on integrative analysis of the data presented in this paper. Color names noted are as per the colors used in Fig. 1D. Region names refer to the ecogeographic groups of accessions based on Sifres et al, 2011. Accession-specific details of mating systems and acylsugar phenotypes are described in Supplementary Figure S9.

One hypothesis for why ASAT4 became inactivated concerns ASAT3, an upstream enzyme in the acylsugar biosynthetic pathway **(Fig. S8)**. This enzyme was studied previously using the same individual plants sampled here for RAD-seq (Schilmiller *et al*., 2015), where it was found that southern accessions had long (>12-carbon) acyl chains on the furanose ring of the sugar molecule **(black squares, Fig. 5)** while this furanose ring acylation was lost in the northern accessions **(red circles, Fig. 5)**. This loss-of-function was associated with loss of the *ASAT3-F* (furanose-ring acylating) duplicate and retention of the *ASAT3-P* (pyranose-ring acylating) duplicate copy of *ASAT3*. Only *ASAT3-P* largely remained in the northern accessions, and this enzyme acylates the same position on the sugar molecule as the ASAT4 enzyme **(Fig. S8)**. These observations led us to hypothesize that ASAT3-P acylation of this site led *ASAT4* to accumulate mutations via drift.

Re-assessed in the context of population structure, the loss of furanose-ring acylation was seen only in the AHZ in clusters 1-5, while clusters 6-8 had acylsugars with furanose-ring acylation. As ASAT3 duplication is predicted to have occurred prior to *S. habrochaites*-*S*.*pennellii* split (Schilmiller *et al*., 2015), presence of active ASAT3-P and -F in cluster 6 (LA1352) provides further support for the origin of *S. habrochaites* near the AHZ southern boundary, where this cluster is located. With the limited data available so far, *ASAT4* inactivation (chemotype E) is seen to have a wider geographic spread than loss of furanose-ring acylation (red circles, **Fig. 5**), suggesting that the two losses occurred independently. However, a deeper genetic and metabolomic sampling of clusters 5, 6 and accessions in cluster 7 close to the southern boundary of the AHZ will need to be performed to better resolve gene loss and gene flow in this geographical area.

## Discussion

In this study, we explored the genetic, reproductive and metabolic diversification of *S. habrochaites* in the context of population structure defined using genome-wide RAD-seq markers. The RAD-seq results provide a less biased estimation of population structure than a previous analysis based on fewer AFLP and SSR markers (Sifres *et al*., 2011), which also tend to be faster evolving than genome-wide SNP loci (Sunde *et al*., 2020). Our results show eight different population clusters across the species range, which have been substantially influenced by variations in geography. The Ecuadorian accessions lie in areas that range from densely forested mountains with high precipitation to either dry or wet coastal regions. On the other hand, accessions south of the AHZ in Peru are often confined to isolated river valleys and experience more uniform environments with regard to altitude and temperature fluctuations. The diverse biotic and abiotic interactions and geographical features are associated with a range of metabolic and reproductive phenotypes in this species (Rick *et al*., 1979; Sacks & St. Clair, 1998; ten Have *et al*., 2007; Finkers *et al*., 2007; Gonzales-Vigil *et al*., 2012; Kim *et al*., 2012; Arms *et al*., 2015; Schilmiller *et al*., 2015; Markova *et al*., 2016; Broz *et al*., 2017b; Kilambi *et al*., 2017; Fan *et al*., 2017), some of which we assessed here.

Previous studies (Rick *et al*., 1979; Sifres *et al*., 2011; Pease *et al*., 2016) have variously placed the origin of *S. habrochaites* in the Huancabamba, Cajamarca or Ancash regions. Here, using findings from multiple genetic analyses and acylsugar genotypes, we infer that the Cajamarca region near the southern boundary of the AHZ is likely the region of *S. habrochaites* origination, supporting Rick *et al*., (1979). Two lineages, representing the two superclusters, then moved north and south from this region to establish the current species range **(Fig. 6)**. Accessions south of the HD are divided into four distinct clusters 5-8. The differentiation between these clusters may have been driven primarily through isolation by distance **(Fig. 3F)**. In contrast, north of the HD, multiple SC populations arose as the species migrated through the AHZ into Ecuador **(Fig. 4B, Table 1)**. Migration through the AHZ and its array of microhabitats may have led to selection for an SC mating system or its fixation due to drift. Limiting mates/pollinators, novel herbivores and/or different temperature/precipitation may have contributed to this evolution. Determining the specific nature of selective pressures experienced in the southern and northern boundaries and in the HD is an interesting research problem that needs to be studied in greater detail. It is noteworthy that the only other tomato clade species on both sides of the AHZ – *S. pimpinellifolium, S. neorickii* – are SC, suggesting that this mating system may be essential for or facilitated by successful migration through the fragmented microhabitats in the AHZ.

As *S. habrochaites* moved through the AHZ, it also accumulated a mutation that led to loss of expression of the *ASAT4* gene involved in acylsugar biosynthesis. Our sampling suggests this expression-inactivated allele is completely restricted to the SI/MP accessions of the AHZ **(Fig. 5; Fig. S9)**. We postulate that this mutation may be an epigenetic modification, since the expression of the allele is seen restored in cluster 2 SC accessions, although *ASAT4* is still inactivated by a new frameshift mutation. It is possible that the association between cluster 2 SC and *ASAT4* inactivation via frameshift is a causative one, in that the emergence of SC in cluster 2 led to rapid fixation of the frameshifted *ASAT4* allele. The fixation of *ASAT4* inactivation in supercluster 1 accessions may also have been accelerated by parallel dynamics occurring at the locus encoding ASAT3, which is upstream in the pathway **(Fig. S8)** creating an epistatic conflict.

Our population structure results and reproductive analyses suggest potential progenitor-descendant relationships between SI and SC populations at both the northern and southern species margins. Given some of the reproductive barriers are uni-directional, there is still potential for continued gene flow between ancestral SI and derivative SC groups. For example, in the case of the SC-2 and the southern SC-4 groups, the loss of pollen SI factors creates a unidirectional inter-population barrier that would prohibit gene flow between SC-2/4 males and progenitor SI females (Markova *et al*., 2016; Broz *et al*., 2017b). In theory, this gene flow could partially rescue SC populations from the evolutionary “dead end” imposed by the mutational loss of SI (Stebbins, 1957; Takebayashi & Morrell, 2001; Igic & Busch, 2013). On the other hand, this partial reproductive barrier can still promote diversification between populations and may represent incipient speciation in *S. habrochaites* at its species margin.

Modern evolutionary synthesis recognizes three modes of speciation: allopatric, parapatric and sympatric, of which allopatric speciation – where reproductive isolation between populations of the same species is driven by geographical barriers (vicariance) – is considered the most common (Allmon, 1992; Howard, 2003). While not serving as a direct cause of vicariant allopatry (Howard, 2003), the AHZ, and especially the HD, appear to have sufficiently destabilized *S. habrochaites* for both reproductive behavior and acylsugar biosynthesis during its northwards migration and set the stage for future reproductive isolation. Evolution of such phenotypic diversity and reproductive isolation sets these populations on a path to forming new species (Allmon, 1992; Howard, 2003). Overall, our findings present a high-resolution view of the micro-evolutionary processes occurring in *S. habrochaites* and provide greater insights into the mechanisms that generate biological diversity.

## Supporting information

SuppFig1

SuppFig2

SuppFig3

SuppFig4

SuppFig5

SuppFig6

SuppFig7

SuppFig8

SuppFig9

## Acknowledgments

We thank the C. M. Rick Tomato Genetics Resource Center for seeds. GM, RL, PAB conceived the study. GM, JL, CM, AKB, AAB, DC, NC, PAB performed the analysis. Everyone contributed to writing and reviewing the manuscript. We thank Dr. A. Tovar-Mendez for providing some of the HT immunoblot results and Nicole Irace for help with pollen tube imaging. This work was supported by grant MCB-1127059 to PAB from the NSF Plant Genome Research Program. JL was supported by the NSF Plant Genome Postdoctoral Fellowship 1711807. GM was supported by IOS-1546617 and IOS-1025636 to RL from the NSF Plant Genome Research Program, and by Cornell University startup funds. AAB was supported by startup funds from Cornell University.

## Supplementary Tables

**Table S1:** TreeMix analysis results. The log likelihood ratio of having zero to ten migration events between the eight coalescent clusters is shown.

| Allowed Migration Events | Log likelihood | LRT (D statistic and p-value) | % variance explained |
| --- | --- | --- | --- |
| 0 | -589.27 | -- | 0.991034 |
| 1 | 57.81 | 1294.15 (2.11e-283) | 0.997558 |
| 2 | 114.99 | 114.37 (1.08e-26) | 0.998156 |
| 3 | 241.72 | 252.46 (4.57e-57) | 0.9992747 |
| 4 | 265.97 | 48.49 (3.32e-12) | 0.9994937 |
| 5 | 285.60 | 39.28 (3.68e-10) | 0.999637 |
| 6 | 302.82 | 34.43 (4.42e-09) | 0.9998275 |
| 7 | 311.70 | 17.78 (2.49e-05) | 0.9999293 |
| 8 | 309.87 | -3.76 (1) | 0.9999032 |
| 9 | 314.70 |  | 0.9999473 |
| 10 | 319.57 |  | 0.9999369 |

## Supplementary Files

**File S1:** Descriptive statistics of accessions, RAD-seq data and their association with other traits assayed in this study. T/B stands for technical or biological replicates.

**File S2:** Primers used for Targeted Sanger sequencing and for S-RNase and HT allele identification

