## Supplementary material for "Migration through a major Andean ecogeographic disruption as a driver of genotypic and phenotypic diversity in a wild tomato species": SuppFig1

### *Figure S1*

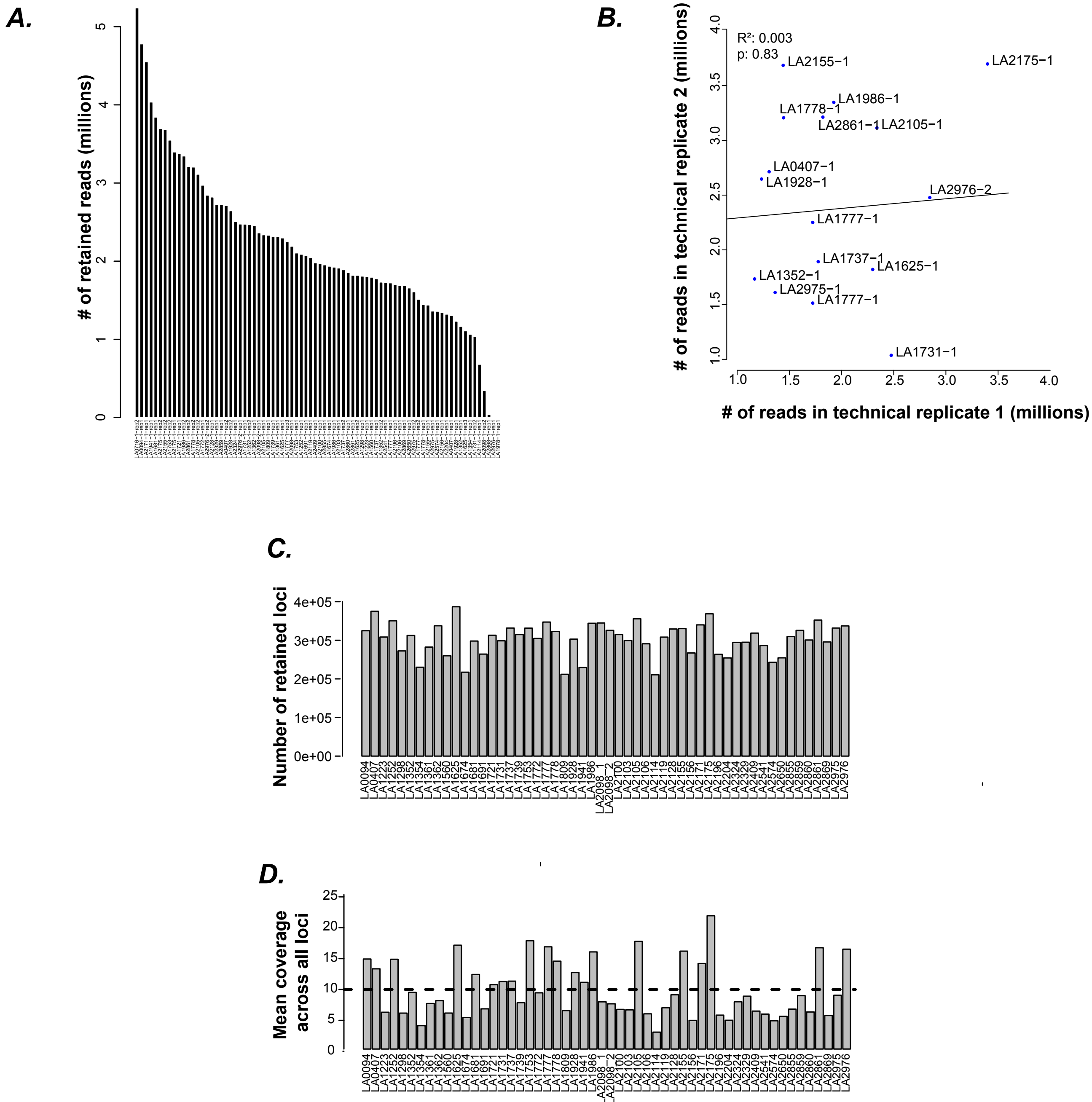


***Supplementary Figure S1: Descriptive statistics of the RAD-seq data.*** (A) # of RAD-seq reads mapped to the LYC4 genome per accession (B) Difference in the number of reads between technical replicates observed is likely due to the randomness of restriction digestion and/or library preparation. -1 and -2 correspond to the first and second biological replicates of the accession. Most accessions had only one biological replicate. All technical replicate samples of the shown accessions were combined together for further analyses. (C) Number of retained loci and (D) Mean read coverage across all loci after running the Stacks pipeline, shown per accession.
