## Supplementary material for "Migration through a major Andean ecogeographic disruption as a driver of genotypic and phenotypic diversity in a wild tomato species": SuppFig2

### *Figure S2*

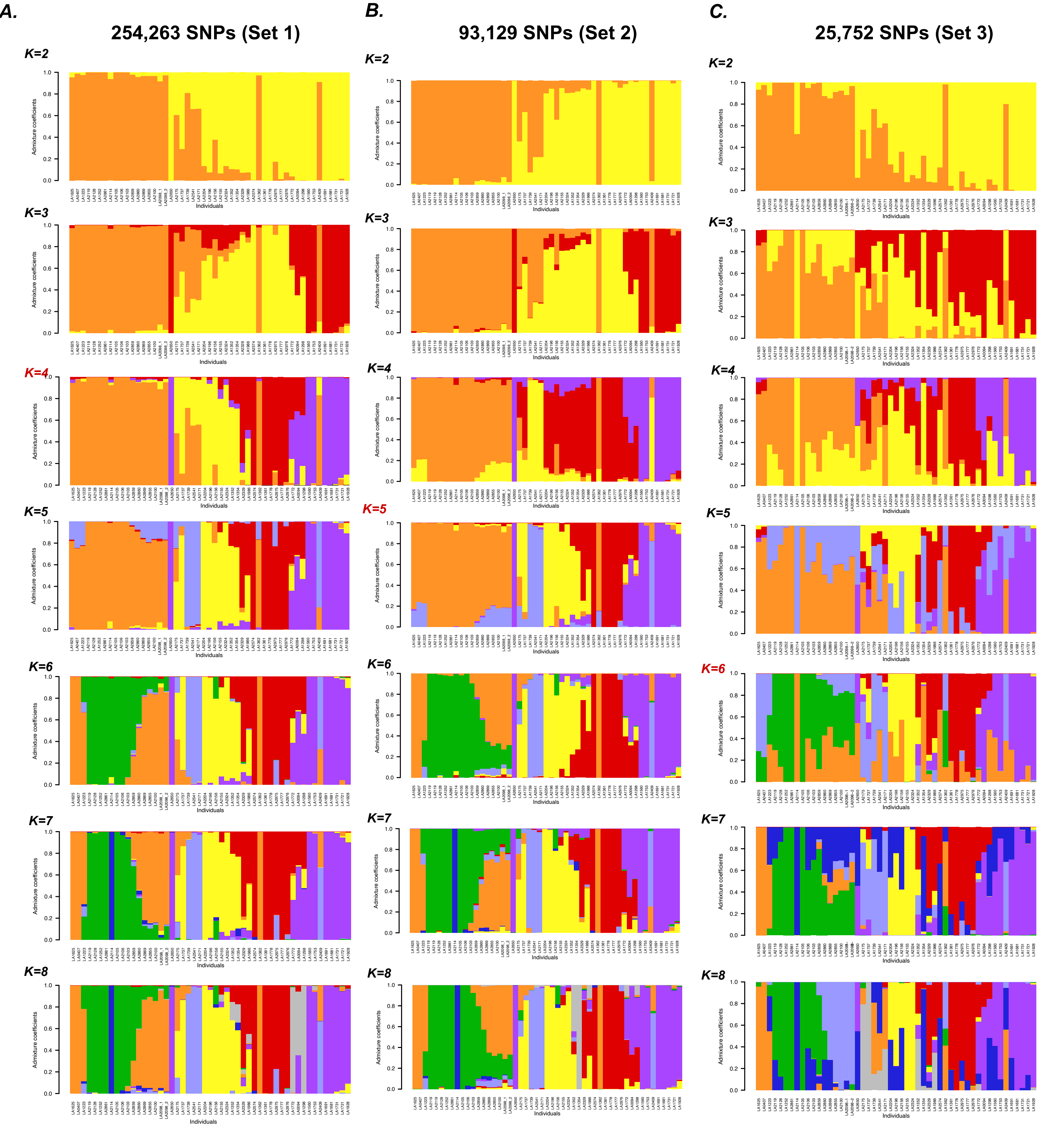


***Supplementary Figure S2: Population structure plots obtained using LEA*** for (A) Set (B) Set 2, (C) Set 3 marker sets. Best K, selected using the cross-entropy criterion, is highlighted in red. There was no substantial change in population assignment after K=6 in all three SNP sets. LA2114 was the only individual to be classified into a different population in K=7, however, the mean read coverage for this accession was the lowest (~2X) and the percentage of missing data the highest (29%, compared to <10% for most other accessions), leading to a lower confidence prediction of the seventh ancestral population.
