## Supplementary material for "Migration through a major Andean ecogeographic disruption as a driver of genotypic and phenotypic diversity in a wild tomato species": SuppFig3

*Figure S3*

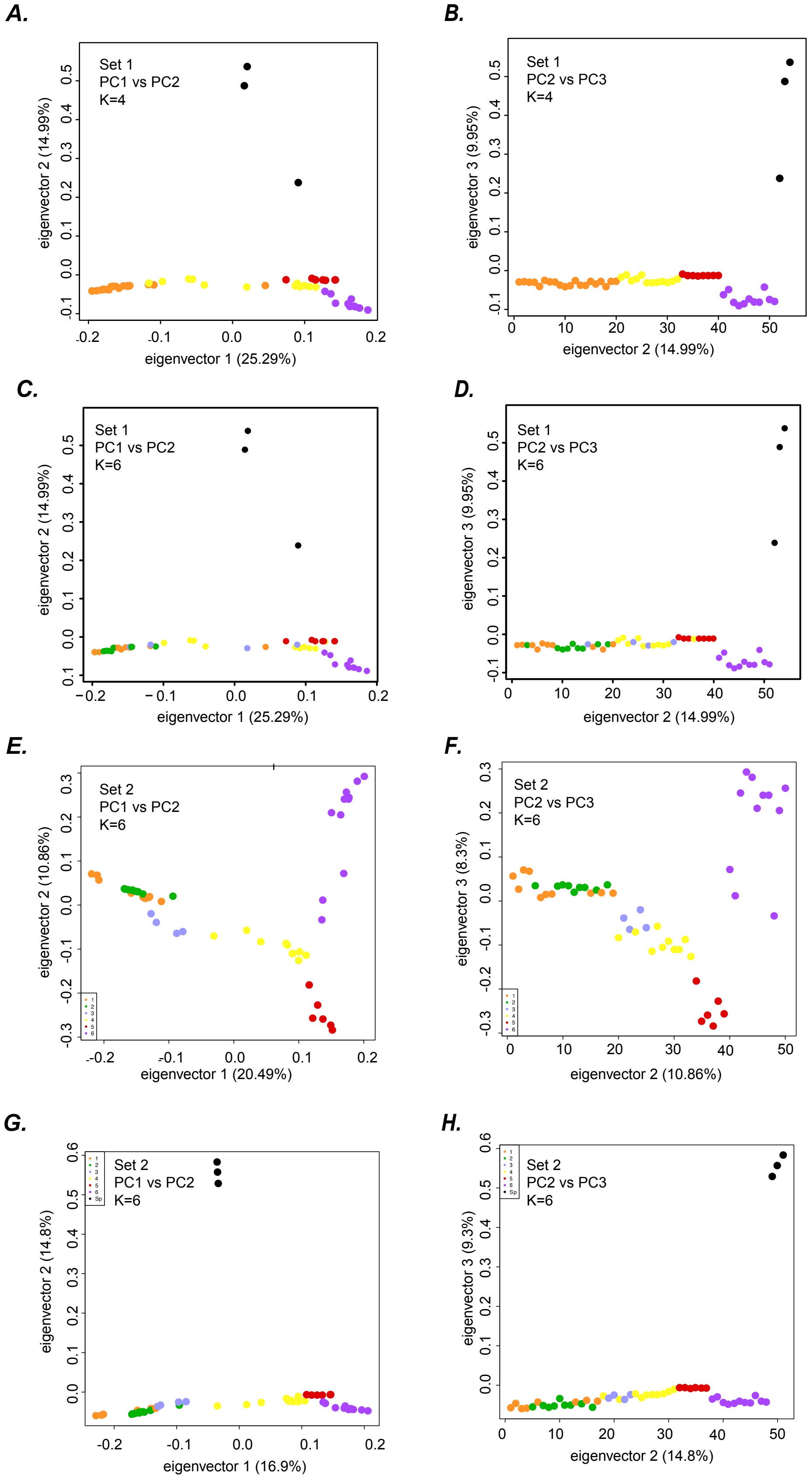


***Supplementary Fig. S3: PCA plots showing population relatedness.*** Different factors tested here for clustering of accessions include # of population clusters (K's), variability explained by the first three PCs, SNP marker sets (Set 1,2), and presence/absence of S. pennellii -- as described in the inset of eac figure.
