## Supplementary material for "Migration through a major Andean ecogeographic disruption as a driver of genotypic and phenotypic diversity in a wild tomato species": SuppFig4

### *Figure S4*

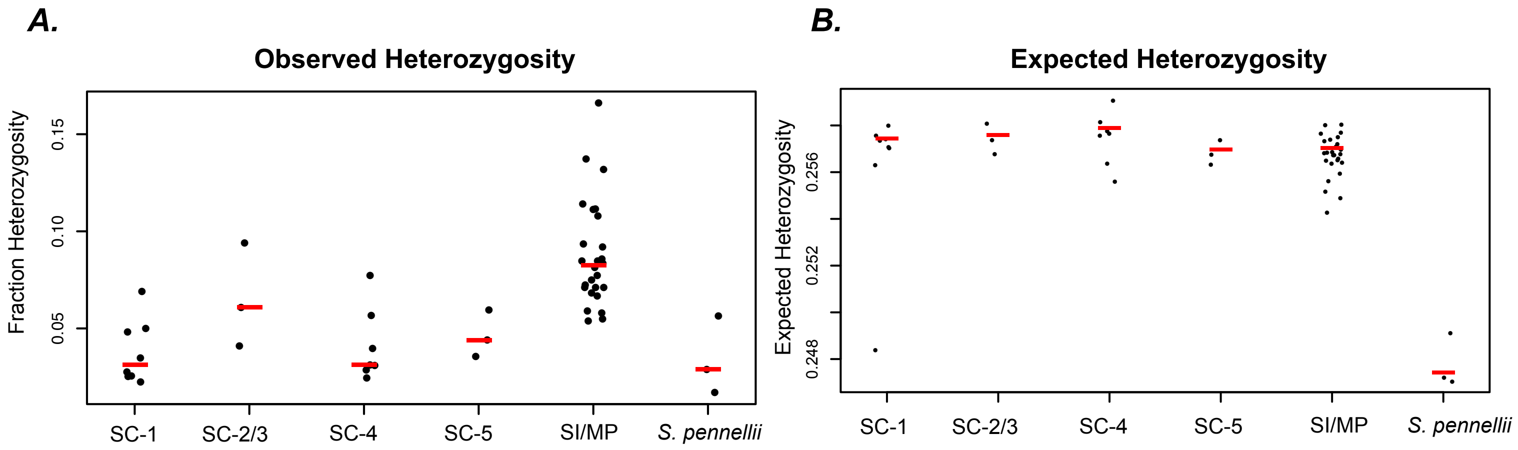


| SC-1 | |
|---|---|
| LA2128 | 0.07 |
| LA2119 | 0.05 |
| LA2114 | 0.05 |
| LA1252 | 0.03 |
| LA2106 | 0.03 |
| LA2103 | 0.03 |
| LA2105 | 0.03 |
| LA2861 | 0.02 |

| SC-2/SC-3 | |
|---|---|
| LA0407 | 0.09 |
| LA1223 | 0.06 |
| LA1625 | 0.04 |

| SC-4 | |
|---|---|
| LA1721 | 0.08 |
| LA1753 | 0.06 |
| LA1731 | 0.04 |
| LA1560 | 0.03 |
| LA1928 | 0.03 |
| LA1691 | 0.03 |
| LA1681 | 0.02 |

| SC-5 | |
|---|---|
| LA2869 | 0.06 |
| LA2860 | 0.04 |
| LA2859 | 0.04 |

| SI/MP | |
|---|---|
| LA2098_1 | 0.17 |
| LA2156 | 0.14 |
| LA1737 | 0.13 |
| LA2175 | 0.11 |
| LA1777 | 0.11 |
| LA1986 | 0.11 |
| LA2976 | 0.11 |
| LA1352 | 0.09 |
| LA1739 | 0.09 |
| LA2098_2 | 0.09 |
| LA2541 | 0.08 |
| LA1298 | 0.08 |
| LA2324 | 0.08 |
| LA1778 | 0.08 |
| LA0094 | 0.08 |
| LA1354 | 0.08 |
| LA1772 | 0.07 |
| LA2100 | 0.07 |
| LA2204 | 0.07 |
| LA2196 | 0.07 |
| LA2155 | 0.07 |
| LA2171 | 0.07 |
| LA1361 | 0.06 |
| LA2329 | 0.06 |
| LA2855 | 0.05 |
| LA2574 | 0.05 |

| *S. pennellii* | |
|---|---|
| LA1674 | 0.06 |
| LA1809 | 0.03 |
| LA1941 | 0.02 |

***Supplementary Fig. S4: Observed and expected heterozygosity divided based on the SC groups as described in the main text.*** Median value is shown using the red line.
