## Supplementary material for "Migration through a major Andean ecogeographic disruption as a driver of genotypic and phenotypic diversity in a wild tomato species": SuppFig5

### *Figure S5*

**A.**

| Accession | # Bags with self fruit, each plant | # Fruits from hand self-pollinations | Pollen tube images (n) | Mating System |
|---|---|---|---|---|
| PI134417 | 3/4, 5/5 | NT | 3 accept, 0 reject (3) | SC |
| PI251305 | 1/3, 1/3 | 2/2, 3/3 | NT | SC |
| PI390515 | 0/3,0/3 | 3/3, 2/2 | 2 accept, 0 reject (2) | SC |
| LA1252 | 3/3, 3/3 | NT | NT | SC |
| LA2128 | 4/5, 2/2 | NT | 2 accept, 0 reject (2) | SC |
| LA2855 | 0/3, 0/4, 2/4 | NT | 1 accept, 6 reject (7) | MP |
| LA2860 | 1/1,1/1 | NT | 5 accept, 0 reject (5) | SC |
| LA4654 | 2/3, 5/7 | NT | NT | SC |
| LA4655 | 3/3 | NT | NT | SC |
| LA4656 | 3/3, 3/3 | NT | NT | SC |

**B.**

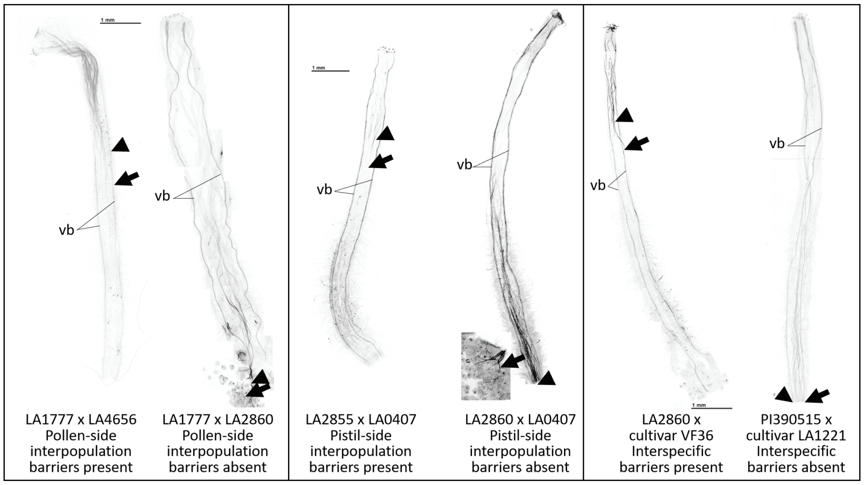


**C.**

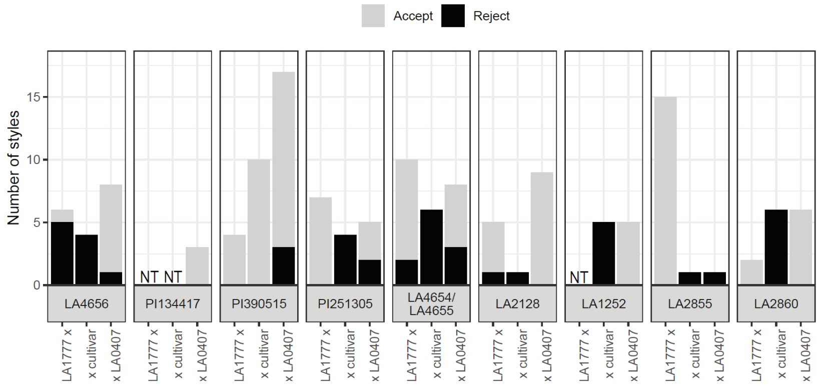


***Supplementary Fig. S5: Definition of mating systems. (A) Determination of mating system for newly phenotyped accessions.*** Inflorescences were covered with net bags to exclude pollinators in order to evaluate whether fruits formed from passive self-pollination, and numbers in the second column indicate the number of bags containing self fruit for different individual. In some cases, hand pollinations were performed to ascertain whether self fruit could be formed, and numbers in the third column indicate the numbers of fruit formed for each hand-pollination for different individuals. In some cases, pollen tube growth in self pollinations was examined, and numbers in the fourth column indicate numbers of images showing pollen tube acceptance (>3 pollen tubes traverse the entire style) or rejection (<3 pollen tubes, typically none, traverse the entire style) with numbers in parentheses (n) indicating the number of individuals tested. The fifth column indicates the final assessment of mating system, SC = self-compatible, MP = mixed population with SC and SI (self-incompatible) individuals. NT = not tested.

***(B) Examples of pollen tube growth analysis for detecting inter-population and interspecific barriers in newly phenotyped accessions.*** Images show pollen tubes in styles stained as described in Materials and Methods. The first pair of images shows crosses with S. habrochaites SI accession LA1777 as the pistil parent. If pollen tubes are rejected, the tested accession exhibits a pollen-side population barrier, here LA4656. If pollen tubes are accepted, this barrier is absent in the tested accession, here LA2860. The second pair of images shows crosses with S. habrochaites SC accession LA0407 as the pollen parent. If pollen tubes are rejected, the tested accession exhibits a pistil-side population barrier, here LA2855. If pollen tubes are not rejected, this barrier is absent in the tested accession, here LA2860. The third pair of images shows crosses with cultivated tomato (S. lycopersicum) as the pollen parent. If pollen tubes are rejected, the tested accession exhibits an interspecific barrier, known as unilateral incompatibility (UI), here LA2860. If pollen tubes are accepted, this barrier is absent in the tested accession, here PI390515. Arrowheads indicate the point at which most pollen tubes stop growing, arrows indicate the longest pollen tube, vb = vascular bundles, which are also stained.

***(C) Histogram summarizing tests for inter-population and interspecific barriers in newly phenotyped accessions.*** Multiple tests were performed as described in Part B for each newly phenotyped accession used in this study. The number of style images showing pollen tube acceptance (>3 pollen tubes traverse the entire style), are shown in gray and images showing pollen tube rejection (<3 pollen tubes traverse the entire style) are shown in black. NT = not tested.
