## Supplementary material for "Migration through a major Andean ecogeographic disruption as a driver of genotypic and phenotypic diversity in a wild tomato species": SuppFig6

### *Figure S6*

```
CLUSTAL multiple sequence alignment by MUSCLE (3.8)

                                                                                                               C1
LA2119_ORF                                   M  Y  K  S  Q  I  L  S  V  L  F  I  S  F  F  A  L  S  T  I  S  I  Y  G  S  F  D  H  W  Q  L  V  L  T  W
S.peruvianum_S13-RNase        ---------------------------------------------------CGTTTTTGCTCTTTCAACCATATCCATTTATGGGTCATTCGACCATTGGCAACTTGTTTTGACTTGGC
LA2119_gDNA                   ---------------------AGTCACAGATCCTGTCAGTTCTCTTCATTTCGTTTTTGCTCTTTCAACCATATCCATTTATGGGTCATTCGACCATTGGCAACTTGTTTTGACTTGGC
LA2119_StylarTranscript       CTTCAAACTAGAGAATGTATAAGTCACAGATCCTGTCAGTTCTCTTCATTTCGTTTTTGCTCTTTCAACCATATCCATTTATGGGTCATTCGACCATTGGCAACTTGTTTTGACTTGGC
                                                                                 ********************************************************************

                                                                 C2
LA2119_ORF                    P  A  G  F  C  K  T  K  D  C  P  R  K  D  I  P  N  N  F  T  I  H  G  V  W  P  D  H  T  S  F  V  M  Y  D  C  D  P  L  K
S.peruvianum_S13-RNase        CAGCAGGCTTCTGCAAAACTAAGGATTGTCCTAGAAAAGATATTCCAAACAACTTTACCATTCATGGCGTTTGGCCAGATCACACCTCCTTCGTCATGTATGACTGTGATCCCCTAAAAA
LA2119_gDNA                   CAGCAGGCTTCTGCAAAACTAAGGATTGTCCTAGAAAAGATATTCCAAACAACTTTACCATTCATGGCGTTTGGCCAGATCACACCTCCTTCGTCATGTATGACTGTGATCCCCTAAAAA
LA2119_StylarTranscript       CAGCAGGCTTCTGCAAAACTAAGGATTGTCCTAGAAAAGATATTCCAAACAACTTTACCATTCATGGCGTTTGGCCAGATCACACCTCCTTCGTCATGTATGACTGTGATCCCCTAAAAA
                              ************************************************************************************************************************

LA2119_ORF                    K  Y  K  T  I  D
S.peruvianum_S13-RNase        AGTATAAAACAATAGAT-------------------------------------------------------------------------------------------------------
LA2119_gDNA                   AGTATAAAACAATAGATGTAAGTGCAAACAGTCATTTTATCAAGCATCTTTTTATATATTTTTTTAAGTTCAACAATTTCCTTCTTGTTTGTTATTTATTTATTTAATCCACTCCTTGCC
LA2119_StylarTranscript       AGTATAAAACAATAGAT-------------------------------------------------------------------------------------------------------
                              *****************

LA2119_ORF                                    D  T  N  I  L  T  E  L  D  A  R  W  P  Q  L  T  S  T  K  I  I  G  L  Q  F  Q  R  F  W  E  Y  E  Y  R  K
S.peruvianum_S13-RNase        ---------------GATACAAATATACTCACTGAACTGGATGCCCGCTGGCCTCAATTGACTTCCACCAAAATAATTGGGTTGCAATTTCAAAGATTCTGGGAATATGAATATAGAAAG
LA2119_gDNA                   TATTGCCTACTACAGGATACAAATATACTCACTGAACTGGATGCCCGCTGGCCTCAATTGACTTCCACCAAAATAATTGGGTTGCAATTTCAAAGATTCTGGGAATATGAATATAGAAAG
LA2119_StylarTranscript       ---------------GATACAAATATACTCACTGAACTGGATGCCCGCTGGCCTCAATTGACTTCCACCAAAATAATTGGGTTGCAATTTCAAAGATTCTGGGAATATGAATATAGAAAG
                                             *********************************************************************************************************

                                      C3                                                  C4
LA2119_ORF                     H  G  T  C  C  A  D  V  F  N  Q  S  M  Y  F  D  I  S  M  K  L  T  D  S  I  D  L  L  K  I  L  R  T  K  G  I  K  P  G  Y
S.peruvianum_S13-RNase        CATGGAACGTGTTGTGCAGATGTCTTCAATCAATCTATGTATTTTGACATATCCATGAAATTAACAGATTCGATTGATCTTTTGAAAATTCTAAGAACAAAGGGAATTAAACCTGGATAT
LA2119_gDNA                   CATGGAACGTGTTGTGCAGATGTCTTCAATCAATCTATGTATTTTGACATATCCATGAAATTAACAGATTCGATTGATCTTTTGAAAATTCTAAGAACAAAGGGAATTAAACCTGGATAT
LA2119_StylarTranscript       CATGGAACGTGTTGTGCAGATGTCTTCAATCAATCTATGTATTTTGACATATCCATGAAATTAACAGATTCGATTGATCTTTTGAAAATTCTAAGAACAAAGGGAATTAAACCTGGATAT
                              ************************************************************************************************************************

                                                                                                                      C5
LA2119_ORF                     T  Y  T  G  D  Q  I  S  R  A  I  K  S  V  T  Q  N  N  P  N  P  K  C  T  Y  I  G  R  S  L  E  L  I  E  I  G  I  C  F  N
S.peruvianum_S13-RNase        ACTTATACCGGTGACCAAATCAGTCGTGCCATTAAGTCAGTTACTCAAAATAATCCAAACCCCAAGTGCACTTATATAGGGAGAAGCTTGGAACTAATTGAGATAGGCATATGTTTCAAC
LA2119_gDNA                   ACTTATACCGGTGACCAAATCAGTCGTGCCATTAAGTCAGTTACTCAAAATAATCCAAACCCCAAGTGCACTTATATAGGGAGAAGCTTGGAACTAATTGAGATAGGCATATGTTTCAAC
LA2119_StylarTranscript       ACTTATACCGGTGACCAAATCAGTCGTGCCATTAAGTCAGTTACTCAAAATAATCCAAACCCCAAGTGCACTTATATAGGGAGAAGCTTGGAACTAATTGAGATAGGCATATGTTTCAAC
                              ************************************************************************************************************************

LA2119_ORF                     R  T  T  N  A  L  M  P  C  P  R  I  S  T  S  C  K  L  G  T  V  E  G  V  K  F  R  *
S.peruvianum_S13-RNase        CGAACAACAAATGCTCTGATGCCATGCCCTCGAATCAGTACGTCATGCAAGTTAGGGACCTTAGAAGGGGTAAAGTTTCGATGATCATCTCTTTCTTTCTTTTTCTTGATAATGTATAAT
LA2119_gDNA                   CGAACAACAAATGCTCTGATGCCATGCCCTCGAATCAGTACGTCATGCAAGTTAGGGACCGTAGAAGGGGTAAAGTTTCGATGATCATCTCTTTCTTTCTTTTTCTTGATAATGTATAAT
LA2119_StylarTranscript       CGAACAACAAATGCTCTGATGCCATGCCCTCGAATCAGTACGTCATGCAAGTTAGGGACCGTAGAAGGGGTAAAGTTTCGATGATCATCTCTTTCTTTCTTTTTCTTGATAATGTATAAT
                              ************************************************************ ***********************************************************

S.peruvianum_S13-RNase        AAAACGTCAGAGACATCCGTGACCGACATAACTATAAAGAAAAATGTTATTCTTGTTGACGGCATTACGGAACCAGTGGAAGGGAATGTCTTGTTAATCAATTTATGATCAGTACAAAAT
LA2119_gDNA                   AAAACG-CAAAGACATCCGTGACCGACATAACTATAAAGAAAAA----------------------------------------------------------------------------
LA2119_StylarTranscript       AAAACG-CAAAGACATCCGTGACCGACATAACTATAAAGAAAAATGTTATTCTTGTTGACGGCATTACGGAACCAGTGGAAGGGAATGTCTTGTTAATCAATTTATGATCAGTACAAAAT

                              ****** ** **********************************

S.peruvianum_S13-RNase        AGCCATACTCAAGATATGAATGTTGGATACCTCGTT
LA2119_gDNA                   ------------------------------------
LA2119_StylarTranscript       AGCCATACTCAAGATATGAATGTTGG----------
```

***Supplementary Fig. S6: The hab-7 S-RNase allele from SC-1 S. habrochaites accession LA2119.*** Alignment of hab-7 S-RNase cDNA and genomic DNA coding sequences is shown (in red), with the intron shown in black. The top line shows the encoded amino acid sequence (ORF). For comparison, alignment with S. peruvianum S13 S-RNase sequences is shown. Conserved S-RNase regions C1-C5 are highlighted.
