## Supplementary material for "Migration through a major Andean ecogeographic disruption as a driver of genotypic and phenotypic diversity in a wild tomato species": SuppFig7

### ***Figure S7***

HT-A genomic sequence alignment

CLUSTAL multiple sequence alignment by MUSCLE (3.8)

```
                M  A  F  K  A  N  I  L  L  I  F  S  L  V  F  M  V  I  S  S  E  V  I  A  R  E  M  V  E
LA0407*         ATGGCATTCAAGGCAAATATTTTGCTTATATTTTCTTTGGTTTTTATGGTTATATCATCAGAGGTTATTGCAAGGGAAATGGTTGAGGGTAAGTTGTTCTAATTGTAGTTTTAAGTACTA
LA4655-1        -----------------TATTTTGCTTATATTTTCTTTGGTTTTTATGGTTATATCATCAGAGGTTATTGCAAGGGAAATGGTTGAGGGTAAGTTGTTCTAATTGTAGTTTTAAGTACTA
LA4655-2        ----------------ATATTTTGCTTATATTTTCTTTGGTTTTTATGGTTATATCATCAGAGGTTATTGCAAGGGAAATGGTTGAGGGTAAGTTGTTCTAATTGTAGTTTTAAGTACTA
LA1223          ATGGCATTCAAGGCAAATATTTTGCTTATATTTTCTTTGGTTTTTATGGTTATATCATCAGAGGTTATTGCAAGGGAAATGGTTGAGGGTAAGTGGTTCTAATTGTAGTTTTAAGTACTA
PI1390515       ATGGCNTTCAAGGCAAATATTTTGCTTATATTTTCTTTGGTTTTTATGGTTATATCATCAGAGGTTATTGCAAGGGAAATGGTTGAGGGTAAGTGGTTCTAATTGTAGTTTTAAGTACTA
PI251305        ATGNCATTCAAGGCAAATATTTTGCTTATATTTTCTTTGGTTTTTATGGTTATATCATCAGAGGTTATTGCAAGGGAAATGGTTGAGGGTAAGTGGTTCTAATTGTAGTTTTAAGTACTA
LA1777*         ATGGCATTCAAGGCAAATATTTTGCTTATATTTTCTTTGGTTTTTATGGTTATATCATCAGAGGTTATTGCAAGGGAAATGGTTGAGGGTAAGTTGTTCTAATTGAAGTTTTAAGTACTA
                                 ***************************************************************************** ********** **************

LA0407*         ATTACAAATTTCATATGACAAATTAAATTAAGTGCATAATGGGATATGGTCAAAGAGAACTTTTACTTTATGAAATATCTTAGTTTTACTCTTTTGTTTTATAGCCTTGTTATTAGCAAA
LA4655-1        ATTACAAATTTCATATGACAAATTAAATTAAGTGCATAATGGGATATGGTCAAAGAGAACTTTTACTTTATGAAATATCTTAGTTTTAC-CCTTTGTTTTATAGCCTTGTTATTAGCAAA
LA4655-2        ATTACAAATTTCATATGACAAATTAAATTAAGTGCATAATGGGATATGGTCAAAGAGAACTTTTACTTTATGAAATATCTTAGTTTTACTCTTTTGTTTTATAGCCTTGTTATTAGCAAA
LA1223          ATTACAAATTTCATATGACAAATTAAATTAAGTGCATAATGGGATATGGTCAAAGAGAACTTTTACTTTATGAAATATCTTAGTTTTAC-CCTTTGTTTTATAGCCTTGTTATTAGCAAA
PI1390515       ATTACAAATTTCATATGACAAATTAAATTAAGTGCATAATGGGATATGGTCAAAGAGAACTTTTACTTTATGAAATATCTTAGTTTTAC-CCTTTGTTTTATAGCCTTGTTATTAGCAAA
PI251305        ATTACAAATTTCATATGACAAATTAAATTAAGTGCATAATGGGATATGGTCAAAGAGAACTTTTACTTTATGAAATATCTTAGTTTTAC-CCTTTGTTTTATAGCCTTGTTATTAGCAAA
LA1777*         ATTACAAATTTCATATCACAAATTAAATTAAGTGCATAATGGCATATGGTCAAAGAGAACTTTTACTTTATGAAATATCTTAGTTTTAC-CCTTTGTTTTATAGCCTTGTTATTAGCAAA
                **************** ************************* ********************************************** * ****************************

LA0407*         ------TCGTCTCATGCAAATGAGTAATTTTGAACCTTTTTTTTTACAAAAGTATTGTAATACGTAAAACTCAACGTCTTTATTGGAGCTCATTTATATTTTAAAATTAAGAGCAAATAT
LA4655-1        ------CCGTCTCATGCAAATGAGTAATTTTGAACCTTTTTTTTACA-AAAGTATTGTAATACGTAAAACTCAACGTCTATATTGAAGCTCATTTAAATTTTAAAATTAAGCGCAAATAT
LA4655-2        ------TCGTCTCATGCAAATGAGTAATTTTGAACCTTTTTTTTTACAAAAGTATTGTAATACGTAAAACTCAACGTCTTTATTGGAGCTCATTTATATTTTAAAATTAAGAGCAAATAT
LA1223          ------TCGTTTCATGCAAATGAGTAATTTTGAACCTTTTTTTTTTACAAAGTATTGTAATATGTAAAACTCAACGTCTATATTGAAGCTCATTTAAATTTTAAAATTAAGCGCAAATAT
PI1390515       ------TCGTTTCATGCAAATGAGTAATTTTGAACCTTTTTTTTTTACAAAGTATTGTAATATGTAAAACTCAACGTCTATATTGAAGCTCATTTAAATTTTAAAATTAAGCGCAAATAT
PI251305        ------TCGTTTCATGCAAATGAGTAATTTTGAACCTTTTTTTTTTACAAAGTATTGTAATATGTAAAACTCAACGTCTATATTGAAGCTCATTTAAATTTTAAAATTAAGCGCAAATAT
LA1777*         TCGTCTTCGTCTCATGCAAATGAGTAATTTTGAAC--TTTTTTTTTACAAAGTATTGTAATACTTAAAACTCAACGTCTATATTGGAGCTCATTTATATTTTAAAATTAAGAGCAAATAT
                       *** ************************  *******    **************  *************** ***** ********** ************** ********

LA0407*         GAGAGAATTTACTAGTGTAAAACTATATAAAATAGACCCAAAATATAATTAAGGGTAAAATTAAGAAATCTTATATAGTAGCATTAGTTGACTTGTCTAAGCAACCATTGATCCCACATG
LA4655-1        GAGAGAATTTACTAGTGTAAAACTATATAAAATAGA-CCAAAATATAATTAAGGGTAAAATTAAGAAATCTTATATAGTAGCATTAGTTGACTTGTCTAAGCAACCATTGATCCCACATG
LA4655-2        GAGAGAATTTACTAGTGTAAAACTATATAAAATAGACCCAAAATATAATTAAGGGTAAAATTAAGAAATCTTATATAGTAGCATTAGTTGACTTGTCTAAGCAACCATTGATCCCACATG
LA1223          GAGAGAATTTACTAGTGTAAAACTATATAAAATAGA-CCAAAATATAATTAAGGGTAAAATTAAGAAATCTTATATAGTAGCATTAGTTGACTTGTCTAAGCAACCATTGATCCCACATG
PI1390515       GAGAGAATTTACTAGTGTAAAACTATATAAAATAGA-CCAAAATATAATTAAGGGTAAAATTAAGAAATCTTATATAGTAGCATTAGTTGACTTGTCTAAGCAACCATTGATCCCACATG
PI251305        GAGAGAATTTACTAGTGTAAAACTATATAAAATAGA-CCAAAATATAATTAAGGGTAAAATTAAGAAATCTTATATAGTAGCATTAGTTGACTTGTCTAAGCAACCATTGATCCCACATG
LA1777*         GAGAGAATTTACCAGTGTAAAACTATATAAAATAGACCCAAAATATAATTAAGGGTAAAATTAAGAAATCTTATATAGTAGCATTAGTTGACTTGTCTAAGCAACCATTGATCCCACATG
                ************ *********************** *********************************************************************************

LA0407*         AAAAAGGTGGTATATTTGACTATATAGAG-AAAAAAAGAAAATTATAATAATTACTATTAAAATTAACATGGAAGAATTAAGAAAATGTTATGAACTTCATTATTATAATTTGATTTTTG
LA4655-1        AAAAAGGTGGTATATTTGACTATATAGAG-AAAAAAAGAAAATTATAATAATTACT---------------------ATGAAGAAAATGTTATGAACTTCATTATTATATTTTGATTTTTG
LA4655-2        AAAAAGGTGGTATATTTGACTATATAGAG-AAAAAAAGAAAATTATAATAATTACTATTAAAATTAACATGGAAGAATTAAGAAAATGTTATGAACTTCATTATTATAATTTGATTTTTG
LA1223          AAAAAGGTGGTATATTTGACTATATAGAGAAAAAAAAGAAAATTATAATAATTACT---------------------ATGAAGAAAATGTTATGAACTTCATTATTATATTTTGATTTTTG
PI1390515       AAAAAGGTGGTATATTTGACTATATAGAGAAAAAAAAGAAAATTATAATAATTACT---------------------ATGAAGAAAATGTTATGAACTTCATTATTATATTTTGATTTTTG
PI251305        AAAAAGGTGGTATATTTGACTATATAGAGAAAAAAAAGAAAATTATAATAATTACT---------------------ATGAAGAAAATGTTATGAACTTCATTATTATATTTTGATTTTTG
```

```
LA1777*      AAAAAGGTGGTATATTTGACTATATAGAGAAAAAAAAGAAAATTATAATAATTACT--------------------ATGAAGAAAATGTTATGAACTTCATTATTATATTTTGATTTTTG
             ***************************** **************************                    ** ***************************** ***********

                                                                         A  N  Q  V  Q  N  S  F  E  L  N  N  P  T  L  Q  K  K  G
LA0407*      AACATATTTCATTTTTTCTGTTCTCTAACAAAATTATTTGTACACAT---CAATTTTGGTGCAGCAAATCAAGTTCAAAATTCATTTGAATTGAATAATCCGACACTTCAGAAAAAAAGGT
LA4655-1     AACATATTTCATTTTTTCTGTTCTCTAACAAAATTATTTGTACACATGACCAATTTTGGTGCAGCAAATCAAGTTCAAAATACATTTGAATTGAATAATCCGACACTTCAGAAAAAAAGGT
LA4655-2     AACATATTTCATTTTTTCTGTTCTCTAACAAAATTATTTGTACACAT---CAATTTTGGTGCAGCAAATCAAGTTCAAAATTCATTTGAATTGAATAATCCGACACTTCAGAAAAAAAGGT
LA1223       AACATATTTCATTTTTTCTGTTCTCTAACAAAATTATTTGTACACATGATCAATTTTGGTGCAGCAAATCAAGTTCAAAATACATTTGAATTGAATAATCCGACACTTCAAAAAAAAAGGT
PI1390515    AACATATTTCATTTTTTCTGTTCTCTAACAAAATTATTTGTACACATGATCAATTTTGGTGCAGCAAATCAAGTTCAAAATACATTTGAATTGAATAATCCGACACTTCAGAAAAAAAGGT
PI251305     AACATATTTCATTTTTTCTGTTCTCTAACAAAATTATTTGTACACATGATCAATTTTGGTGCAGCAAATCAAGTTCAAAATACATTTGAATTGAATAATCCGACACTTCAGAAAAAAAGGT
LA1777*      AACATATTTCATTTTTTCTGTTCTCTAACAAAATTATTTGTACACAT---CAATTTTGGTGCAGAAAATCAAGTTCAAAATACATTTGAATTGAATAATCCGACACTTCAGAAAAAAAGGT
             ***********************************************   ************** *************** ************************** ********

              G  G  S  L  F  P  N  I  A  C  L  G  C  S  C  P  K  K  D  N  K  N  N  N  N  N  N  N  N  N  D  D  D  D  D  D  D  S  F  I
LA0407*      GGGGGATCATTATTTCCTAATATAGCGTGTTTGGGTTGCAGTTGCCCAAAAAAAGATAATAAAAACAATAATAATAATAATAATAATAACGATGACGATGATGACGATGATAGTTTCATT
LA4655-1     GGGGGATCATTATTTCCTAATATAGCGTGTTTGGGTTGCAGTTGCCCAAAAAAAGATAATAAAAACAATAATAATAATAATAATAATAACGATGACGATGATGACGATGATAG-------
LA4655-2     GGGGGATCATTATTTCCTAATATAGCGTGTTTGGGTTGCAGTTGCCCAAAAAAAGATAATAAAAACAATAATAATAATAATAATAATAACGATGACGATGATGACGATGATAG-------
LA1223       GGGGGATCATTATTTCCTAATATAGCGTGTTTGGGTTGCAGTTGCCCAAAATAAGATAATAAAAAC------AATAATAATAATAATAACGATGACGATGATGACGATGACAGTTTCATT
PI1390515    GGGGGATCATTATTTCCTAATATAGCGTGTTTGGGTTGCAGTTGCCCAAAATAAGATAATAAAAAC------AATAATAATAATAATAACGATGACGATGATGACGATGACAG-------
PI251305     GGGGGATCATTATTTCCTAATATAGCGTGTTTGGGTTGCAGTTGCCCAAAATAAGATAATAAAAAC------AATAATAATAATAATAACGATGACGATGATGACGATGA----------
LA1777*      GGGGGATCATTATTTCCTAATATAGCGTGTTTGGGTTGCAGTTGCCCAAAAAAAGATAATAAAAAC------AATAATAATAATAATAACGATGACGATGATGACGATGACAGTTTCATT
             *************************************************** *************      *************************************

              G  N  V  C  K  A  M  C  C  *
LA0407*      GGTAATGTTTGTAAAGCCATGTGTTGTTAG
LA4655-1     ------------------------------
LA4655-2     ------------------------------
LA1223       GGTAATGTTTGTAAAGCCATGTGTTGTTAG
PI1390515    ------------------------------
PI251305     ------------------------------
LA1777*      GGTAATGTTTGTAAAGCCATGTGTTGTTAG
```

***Supplementary Fig. 7: Alignment of HT-A genomic DNA sequences.*** HT-A sequences were amplified using gene specific primers and PCR products (LA4655, PI1251305, PI1390515) or clones of PCR products (LA1223) were subject to Sanger sequencing. Sequences were aligned with references from LA0407 (Genbank GU362659.1) and LA1777 (GU362649.1). Exons are displayed in red, and the deduced amino acid sequence is denoted above the exons. The A->T SNP leading to a nonsense mutation (K->stop) in exon 2 that was discovered in some northern accessions is highlighted in gray. *Covey et al. 2010.
