## Supplementary material for "Migration through a major Andean ecogeographic disruption as a driver of genotypic and phenotypic diversity in a wild tomato species": SuppFig8

### *Supplementary Figure S8*

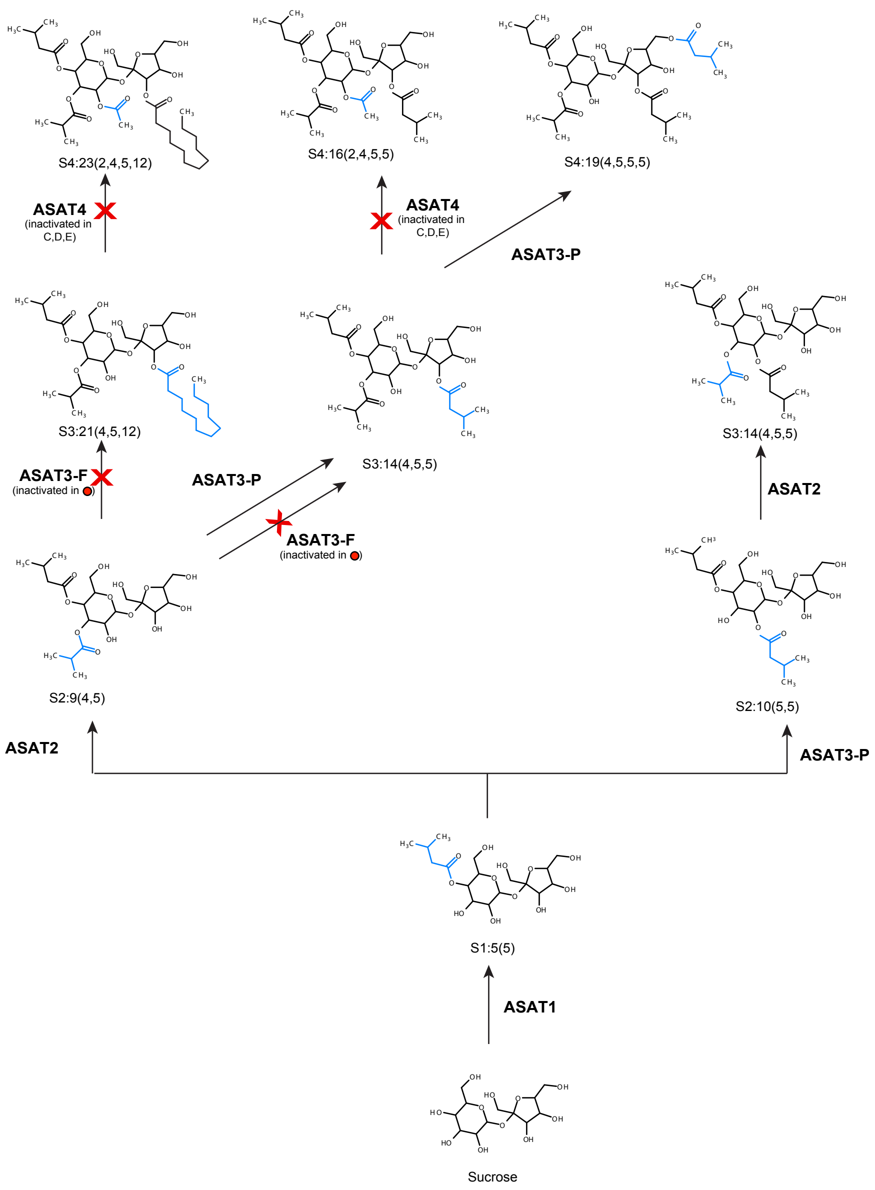


***Supplementary Figure S8: Acylsugar diversity in S. habrochaites is a result of diverse promiscuous activities of the acylsugar acyltransferases (ASATs).*** The trichome acylsugar biosynthetic pathway in *S. habrochaites* is shown. ASAT3-F and ASAT4 loss patterns shown in Fig. 5 can be understood in the context of the pathway shown here. Grey colored ASAT3-P step incicates enzyme is present but substrate is not made due to upstream inactivations. * *ShASAT3-F* sequence is not absent in all E-chemotype accessions (e.g. LA1352).Each enzyme has preferences for using certain types of CoAs. These preferences can be found in Schilmiller et al, Plant Cell, 2015 and Fan et al, Current Opinion in Plant Biology, 2019. Activities shown for ASAT2 are from ASAT2-FP (LA1777), as described in Fan et al, Current Opinion in Plant Biology, 2019.
