## Supplementary material for "Migration through a major Andean ecogeographic disruption as a driver of genotypic and phenotypic diversity in a wild tomato species": SuppFig9

*Figure S9*

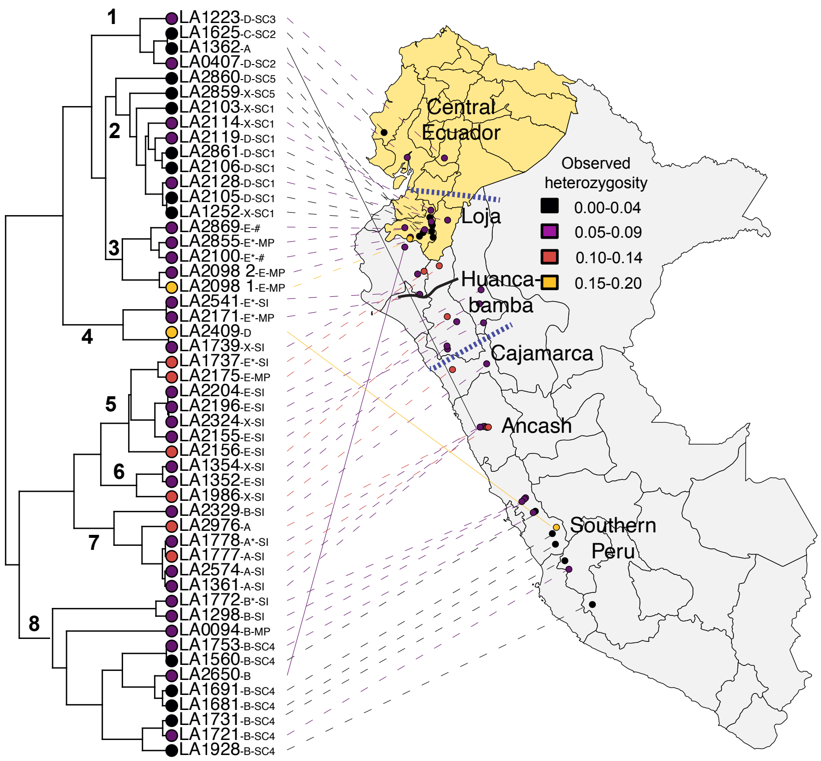


***Supplementary Figure S9: Summary of phenotypes measured for S. habrochaites.*** Heterozygosity estimates are associated with the phylogenetic tree obtained using coalescent analysis. Yellow highlighted regions corresponds to Ecuador, while the rest of the map represents Peru. The AHZ spans the region bounded by the blue lines, with the Huancabamba Depression indicated by the solid black curve. Named regions are as per Sifres et al, 2011. Letters after accession numbers stand for acylsugar chemotype based on Kim et al, 2012 and Fig. 5A (if marked by *), followed by SC/SI/MP assignment as per Fig. 4A. X=no chemotype assignment, #=mating system not assessed.
